# Bibacillin 1: A two-component lantibiotic from *Bacillus thuringiensis*

**DOI:** 10.1101/2024.08.13.607848

**Authors:** Ryan Moreira, Yi Yang, Youran Luo, Michael S. Gilmore, Wilfred van der Donk

## Abstract

Here we describe bibacillin 1 – a two-component lantibiotic from *Bacillus thuringiensis*. The peptides that comprise bibacillin 1 are modified by a class II lanthipeptide synthetase Bib1M producing two peptides with non-overlapping ring patterns that are reminiscent of cerecidin and the short component of the enterococcal cytolysin (CylLS”), a virulence factor associated with human disease. Stereochemical analysis demonstrated that each component contains LL-methyllanthionine and DL-lanthionine. The mature bibacillin 1 peptides showed cooperative bactericidal activity against Gram-positive bacteria, including members of ESKAPE pathogens, and weak hemolytic activity. Optimal ratio studies suggest that bibacillin 1 works best when the components are present in a 1:1 ratio, but near optimal activity was observed at ratios strongly favouring one component over the other, suggesting that the two peptides may have different but complementary targets. Mechanism of action studies suggest a lipid II-independent killing action distinguishing bibacillin 1 from two other two-component lantibiotics haloduracin and lacticin 3147. One of the two components of bibacillin 1 showed cross reactivity with the cytolysin regulatory system. These result support the involvement of bibacillin 1 in quorum sensing and raise questions about the impact of CylL_S_”-like natural products on lanthipeptide expression in diverse bacterial communities.

## Introduction

Antibiotic resistance continues to pose a global health risk as morbidity and mortality due to drug resistant bacteria increases. The economic impact of antibiotic resistance carries a cost of tens of billions of dollars in the United States of America each year.^1^ A wide variety of challenges must be overcome to address this problem.^2^ One important step is the discovery of new antibiotics, preferably compounds with unique action mechanisms that circumvent cross-resistance. Natural products are a proven source for antibiotics.^3,4^ Recently, a renewed interest in natural products has emerged due to advances in DNA sequencing and genome mining that has provided new avenues by which new natural product antibiotics can be discovered.^5^ Part of this renaissance are ribosomally synthesized and post translationally modified peptides (RiPPs). In addition to their diverse bioactivities,^6^ RiPPs are a particularly interesting class of natural products because substrates and modifying enzymes are often encoded within the same biosynthetic gene cluster (BGC) making genome mining using bioinformatic tools and heterologous expression relatively simple.^7^

An important and antibiotic-rich class of RiPPs are the lanthipeptides, which are peptide natural products that contain lanthionine bridges.^6,8^ These thioether cross links are installed enzymatically through dehydration of Ser and Thr residues giving dehydroalanine (Dha) and dehydrobutyrine (Dhb), respectively, followed by enzyme-guided Michael-type addition by the Cys sulfhydryl group to form lanthionine (Lan) and methyllanthionine (MeLan), respectively. The resulting bridges rigidify the peptide conferring defined secondary structure and imbuing this class of natural products with a range of attractive properties.^6,9–11^ When a lanthipeptide possesses antibiotic activity, it is often referred to as a lantibiotic.^12^ Nisin is the best characterized lantibiotic since it has been an FDA approved food preservative for nearly four decades.^13^ Another important prototype lantibiotic is mersacidin, which possesses an overlapping ring pattern giving it a globular structure.^14^ Despite their structural differences, both nisin and mersacidin target the essential cell wall precursor lipid II.^15–18^

Two-component lantibiotics are an enigmatic subclass of lantibiotics.^19^ These systems consist of two structurally distinct peptides that work together to kill bacterial cells. The best understood two-component lantibiotics are haloduracin and lacticin 3147.^20–23^ Both of these lantibiotics possess a mersacidin-like component that binds to lipid II (the α peptide) and an extended component that initiates pore formation when it binds to the peptide:lipid II complex (the β peptide).^22,24–27^ A few two-component lantibiotics are known that do not possess a mersacidin-like component.^28–30^ The best known is the enterococcal cytolysin composed of a short subunit CylL_S_” and long subunit CylL_L_” that both possess non-overlapping ring patterns (Fig. 1A).^31^ The unique structure of cytolysin gives it remarkable bioactivity as it is the only known lantibiotic that targets both Gram-positive bacteria and eukaryotic cells.^28^ Enterococcal cytolysin is a hemolytic virulence factor linked to human disease and increased patient mortality.^32,33^

**Fig. 1.**
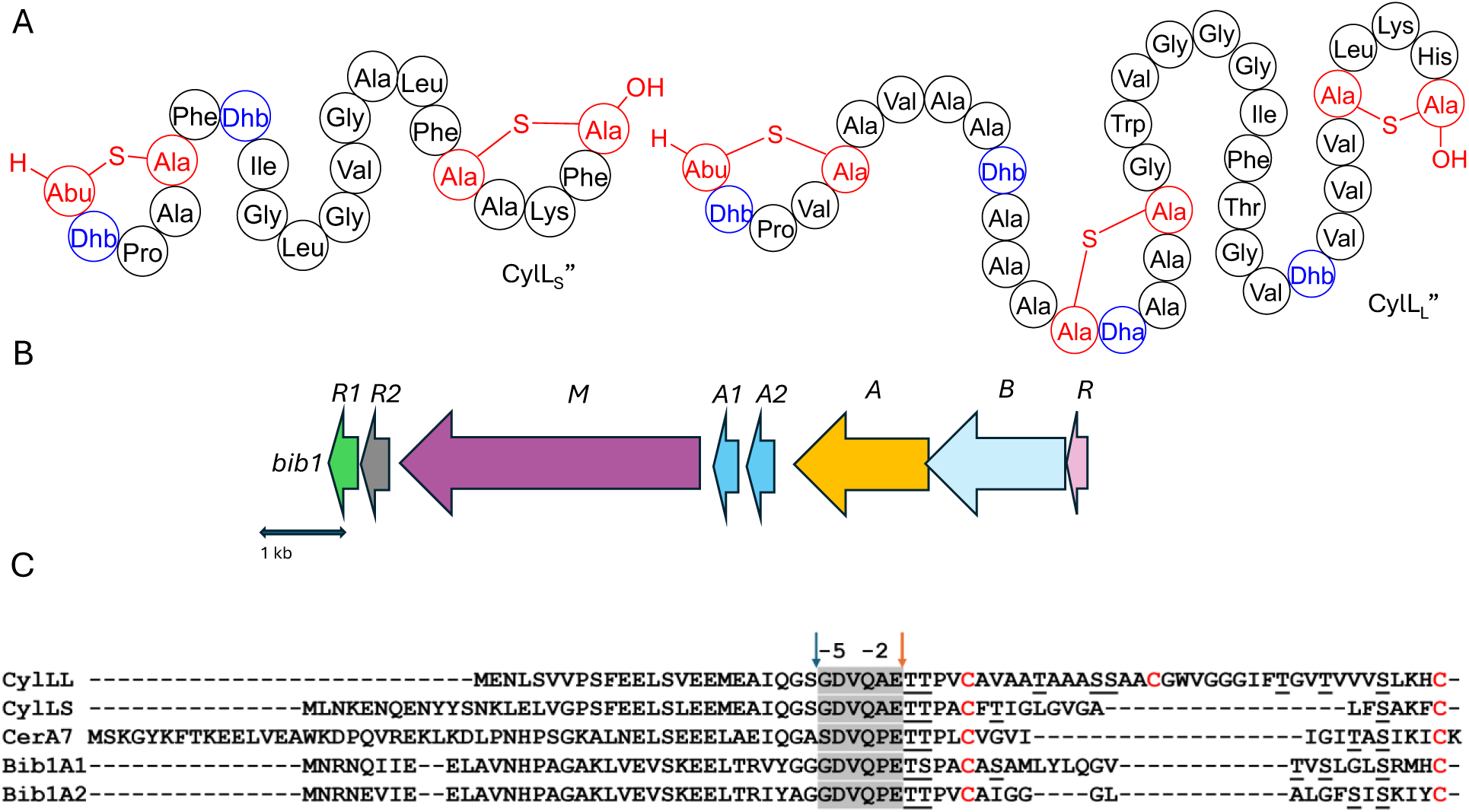
A) The structures of CylL_L_” and CylL_S_”. Residues involved in Lan/MeLan are colored red and dehydrated residues are colored blue. B) The *bib1* biosynthetic gene cluster. C) Sequences related to Bib1A1 and Bib1A2. Blue and orange arrows indicate protease cleavage sites during the maturation of CylL_S_” and CylL_L_” by CylB and CylA, respectively. The hexapeptide sequence removed during the second proteolytic cleavage is highlighted in light grey. Cysteine residues are highlighted in red and dehydratable residues are underlined.

In previous work, while exploring the evolutionary relationship between the single peptide lantibiotic cerecidin^34^ and the two-component system cytolysin we uncovered many BGCs encoding cytolysin-like peptides.^35^ A subset of these BGCs encode two distinct peptides of different lengths that differ from cytolysin suggesting that they constitute undescribed two-component lantibiotics. These peptides were named bibacillin I when both peptides were predicted to contain two thioether rings, and bibacillin II when one of the peptides contains only one thioether ring. At the time of the discovery of their BGCs, they were found mainly in the bacillus genera. Because of the unique structures of these peptides, which suggest they may have a distinct mechanism of action, we aimed to characterize an example. Here, we elucidate the structure of a representative bibacillin I class member, bibacillin 1 (Bib1) from *Bacillus thuringiensis*, and characterize its bioactivity. A preliminary investigation of action mechanism and quorum sensing activity is also presented.

## Results and discussion

### Establishing the structures of Bib1α and Bib1β

Assessment of the genes surrounding those that encode the Bib1 substrate peptides using the program RODEO^36^ revealed a biosynthetic architecture consistent with class II lanthipeptides.^37^ Promoter prediction tools^38^ were used to narrow the selection of open reading frames from the RODEO output to only include those of biosynthetic relevance (Fig. 1B). When naming genes in the Bib1 cluster, we opted to mostly use annotations that are analogous to those found in the cytolysin cluster (*cyl*).^39,40^ The putative BGC encodes Bib1M, a class II synthetase bearing a resemblance to CylM^41^ (48.7 % similarity). In addition, the cluster encodes a LuxR-like sequence, termed Bib1R that bears a strong resemblance to CerR (65.5% similarity), which is known to regulate production of and protect *Bacillus cereus* from cerecidins.^42^ Adjacent to the gene for Bib1M, the BGC contains two genes encoding proteins that are similar to the cytolysin quorum sensing system CylR1 and CylR2,^43^ named here Bib1R1 and Bib1R2, suggesting that Bib1 might also serve a role in quorum sensing. We deviate from the nomenclature of the cytolysin BGC when naming the precursor peptides and genes and instead use terminology previously introduced in the lanthipeptide field.^44^ We termed the slightly longer peptide Bib1A1 and the slightly shorter peptide Bib1A2 (Fig. 1C).

RiPP precursors are composed of a core peptide, which undergoes post-translational modification and comprises the mature natural product, and a leader peptide, which does not undergo post-translational modification, but is necessary for substrate recognition by biosynthetic enzymes during maturation and controls product toxicity.^8,45^ Therefore, an important step when biosynthesizing RiPPs is determining where to cleave the modified substrate to remove the leader peptide from the mature natural product. The primary sequences of Bib1A1 and Bib1A2 are consistent with class II lanthipeptides and are similar to those of the precursors to cytolysin (Fig. 1C). During maturation, cytolysin undergoes two proteolytic cleavages. First, the bifunctional protease-transporter CylB cleaves modified CylL_L_ (mCylL_L_) and mCylL_S_ at a double glycine motif (GG, GS, AG, or GA, Fig. 1C), and secretes the C-terminal modified core peptide.^39,46^ CylA, an extracellular serine protease, then removes what remains of the leader peptide (six amino acids, Fig. 1C) producing the fully mature CylL_L_” and CylL_S_”.^46,47^ Considering the homology between CylB and Bib1B, and CylA and Bib1A, we suspected that Bib1A1 and Bib1A2 would undergo a similar sequential proteolysis to achieve maturity. Bib1A1 and Bib1A2 contain a unique Pro in the C-terminal hexapeptide of the leader sequence that differs from the Ala residue found in the corresponding cytolysin hexapeptides (Fig. 1C). Previous studies showed that CylA can remove the entire leader peptide in one step in vitro.^35,48^ To allow for removal of the leader peptides using CylA after co-expression of precursor peptides with Bib1M, the hexapeptide sequence was mutated at a single position to match the cytolysin sequence (Pro−2Ala). Henceforth, Bib1A1 and Bib1A2 refer in this report to the Pro−2Ala mutants of these substrate peptides.

We chose to access bibacillin 1 by heterologous production in *Escherichia coli*. Many lanthipeptides have been successfully produced in this heterologous host, with class II lanthipeptides particularly amenable.^29,34,49–59^ In all cases where the natural product was known, the product in *E. coli* had the same structure (ring pattern, stereochemistry, dehydration extent). Co-expression of Bib1M with His_6_-Bib1A1 in *E. coli* followed by immobilized metal affinity chromatography (IMAC) yielded a modified peptide that had undergone six dehydrations. Removal of the leader peptide with CylA occurred smoothly giving the mature product of Bib1A1 that we call Bib1α. HPLC purification followed by high resolution LCMS-MS analysis revealed that Bib1α possessed a non-overlapping ring pattern similar to the CylL_S_” ring pattern (Fig. 2A,C). Co-expression of His_6_-Bib1A2 with Bib1M yielded a peptide that had undergone three dehydrations. As with Bib1A1, leader peptide removal with CylA went smoothly allowing for the isolation of Bib1β. Analysis of this peptide via high resolution LCMS-MS revealed that it also possessed a CylL_S_”-like ring pattern (Fig. 2B,C).

**Fig. 2.**
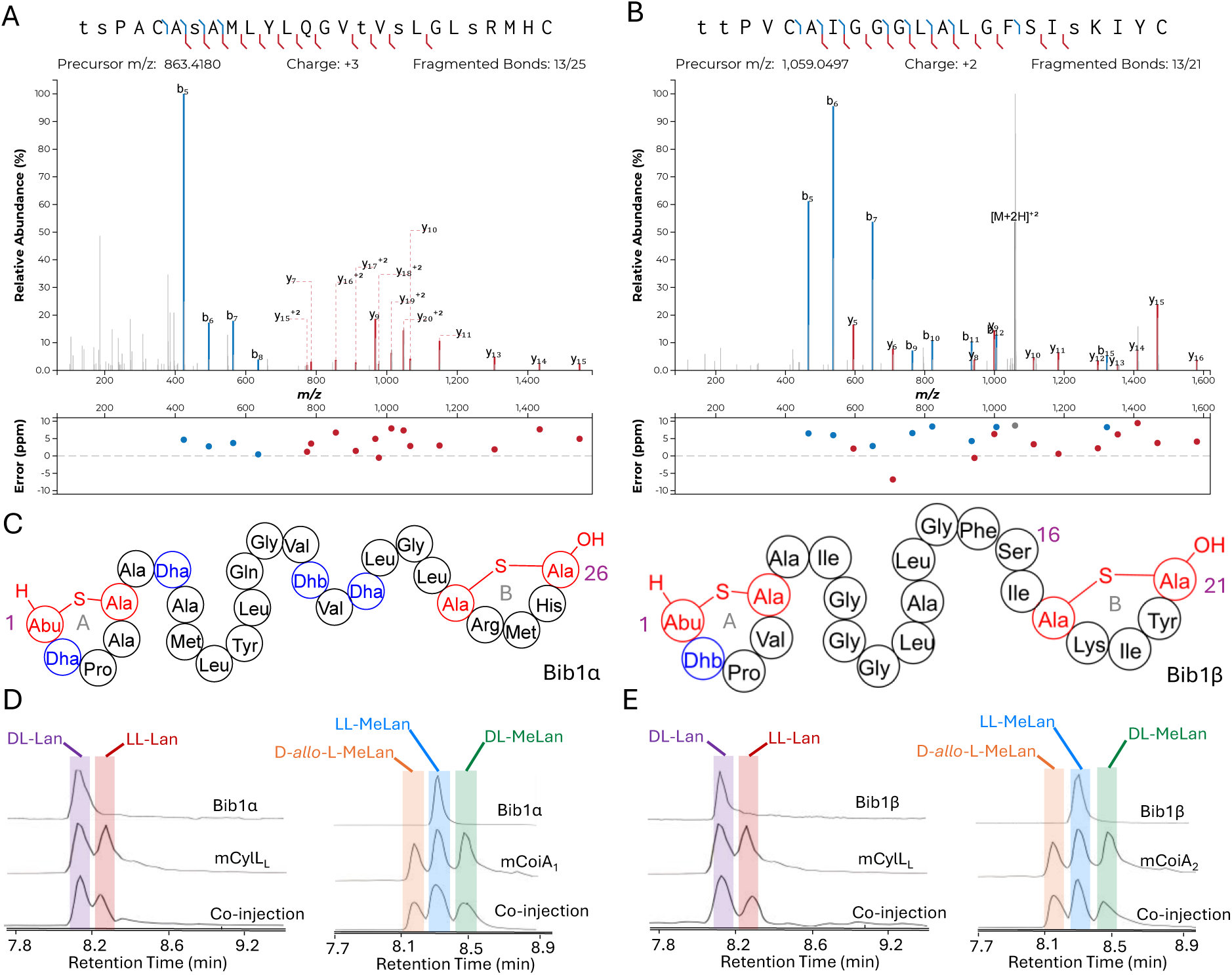
Structural characterization of Bib1α and Bib1β. A, B) MS-MS data of Bib1α and β, respectively. Plots were prepared using the Interactive Peptide Annotator Webtool.^60^ Colored lines denote b and y ions and lower case letters denote dehydrated residues. Differences between theoretical masses and the masses detected are presented in a graph below each spectrum. C) Ring patterns of Bib1α and Bib1β that are consistent with the MS-MS data. D, E) Marfey’s analysis of the Lan and MeLan bridges in Bib1α and Bib1β, respectively. For analysis of Lan, mCylL_L_ was used as a standard. For analysis of MeLan, mCoiA_1_ was used as a standard.

The structure of Bib1α and Bib1β have similarities and differences to CylL_S_”. Overall, all three peptides are hydrophobic and slightly cationic. The A and B rings found in Bib1α, Bib1β and CylL_S_” are the same size, and each peptide contains an A ring formed from a Dhx-Dhx-Xxx-Xxx-Cys motif (Dhx refers to Dha or Dhb). Bib1α contains five more residues than CylL_S_” which extends the distance between the A and B rings. In contrast, Bib1β contains only one more residue than CylL_S_”. CylL_S_” and Bib1α have only hydrophobic residues in the sequences connecting their A and B rings including dehydroamino acids, but Bib1β contains a Ser at position 16 and contains no dehydroamino acids in the sequence between the rings. Like CylL_S_” and CylL_L_”, both bibacillin peptides possess a positively charged residue in the B ring (Arg in Bib1α and Lys in Bib1β), but Bib1α and Bib1β are more similar to CylL_S_” than CylL_L_”.

Analysis of the absolute configuration of the Lan and MeLan residues in Bib1α and Bib1β was conducted using an optimized protocol recently described that employs modified Marfey’s reagents and Lan/MeLan standards of known configuration that are prepared biosynthetically.^58^ Using this approach, we determined that Bib1α and Bib1β both possess one LL-MeLan and one DL-Lan (Fig. 2D-E) further establishing their similarity to CylL_S_” (Fig. 1A).^31^ The stereochemistry of the MeLan residues found in bibacillin 1 is consistent with previous studies showing that Dhx-Dhx-Xxx-Xxx-Cys sequences favour the formation of LL-MeLan through substrate templating.^54^ Haloduracin α and both peptides of cytolysin possess this N-terminal sequence motif and all three possess LL-MeLan.

Class II synthetases, like Bib1M, are known to dehydrate Ser/Thr residues through ATP-driven phosphorylation followed by elimination of the phosphate group.^41^ When co-expressing class II synthetases with their cognates substrate(s) it is not uncommon to find small amounts of phosphorylated peptide. Recently, the phosphorylated form of a lanthipeptide was found to be the mature natural product.^11^ Approximately 30-50% of His_6_-Bib1A2 was phosphorylated at Ser64 (Ser16 in the mature natural product) during co-expression with Bib1M in *E. coli*, as indicated by MS-MS characterization (Fig. S1, ESI†), raising the question of whether the mature peptide may be unmodified, phosphorylated, or dehydrated at this position. To assess all three possibilities, we isolated the phosphorylated form of Bib1β (which we termed Bib1β_P_) and devised a semi-synthetic approach to the fully dehydrated form of Bib1β (named Bib1β_Dha_). Cys-containing proteins and peptides can be converted to Dha-containing products through chemical mutagenesis.^61,62^ Analogously, we envisioned preparing Bib1β_Dha_ by replacing the partially phosphorylated Ser16 with a Cys residue, and then chemically converting this residue to a Dha (Fig. 3A). Treatment of Bib1M-modified His_6_-Bib1A2-S64C with 2,5-dibromovalerate/K_2_CO_3_ converted this peptide to the desired His_6_-Bib1A2-S64Dha.^63^ Buffer exchange followed by treatment of modified His_6_-Bib1A2-S64Dha with CylA yielded Bib1β_Dha_. Analysis of Bib1β_Dha_ via high resolution LCMS-MS revealed that it possessed the same ring pattern as Bib1β and a Dha at position 16 (Fig. 3B).

**Figure 3.**
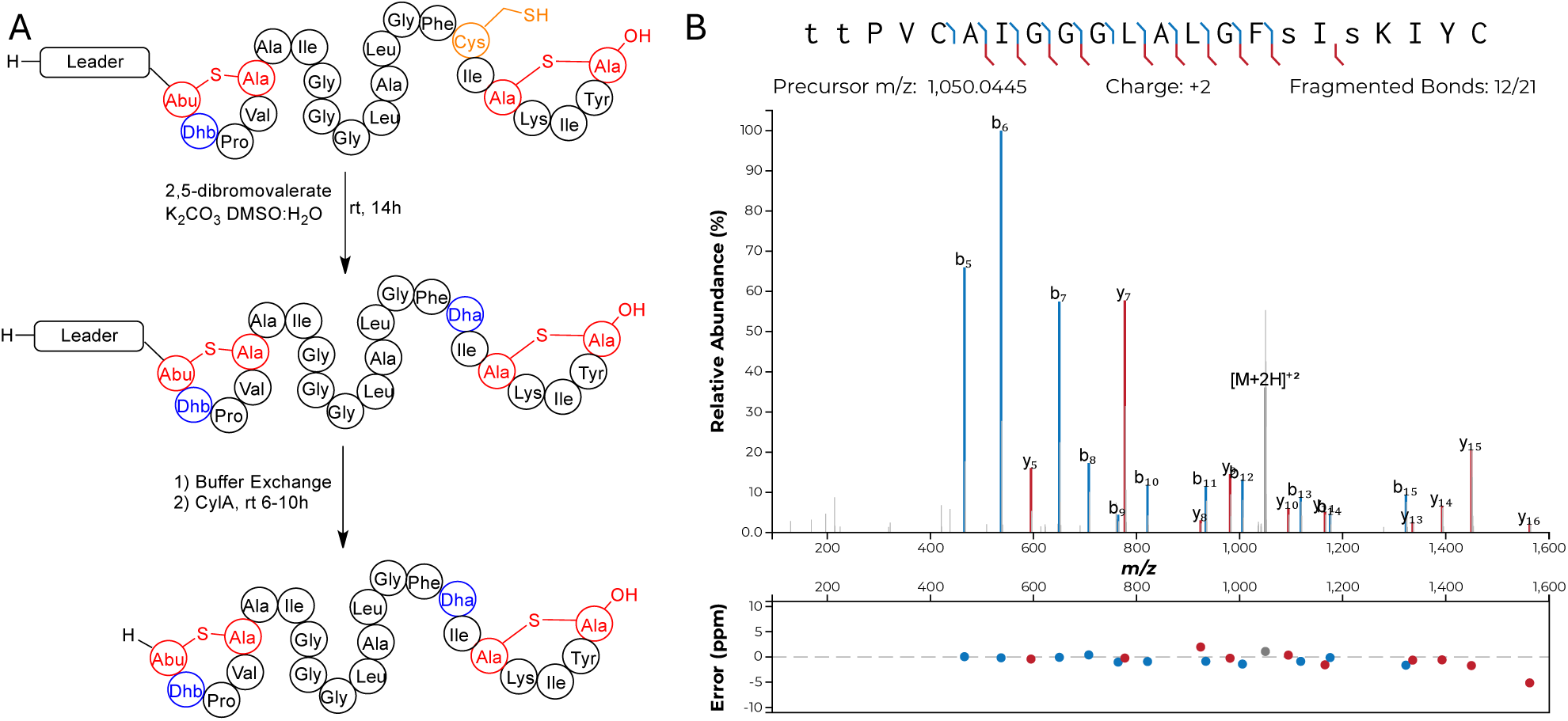
Synthesis of Bib1β_Dha_ through chemical mutagenesis. A) Synthetic scheme showing the conversion of modified His_6_-Bib1A2-S64C to Bib1β_Dha_. B) MS-MS analysis of Bib1β_Dha_ demonstrating that its ring pattern is the same as WT Bib1β. Colored lines denote b and y ions and lower-case letters denote dehydrated residues. Differences between theoretical masses and the masses detected are presented in a graph below each spectrum.

### Antimicrobial and hemolytic activity

Considering the structural similarity to CylL_S_” we wondered if bibacillin 1 would display similar bioactivity compared to cytolysin. We first analyzed the hemolytic activity of bibacillin 1. On an agar plate, Bib1α and β showed hemolytic activity when administered individually or in a 1:1 ratio (Fig. 4A). Bib1β_P_ or Bib1β_Dha_ did not show any hemolytic activity beyond that of a negative control but did show weak hemolytic activity when combined with Bib1α in a 1:1 ratio. A solution phase assay demonstrated Bib1α and Bib1β did not induce hemolysis at concentrations of 2 µM, even when administered in a 1:1 ratio (Fig. S2A, ESI†). In contrast, CylL_L_”:CylL_S_” readily lysed rabbit erythrocytes with an EC_50_ of 76 ± 5 nM when administered in a 1:1 ratio (Fig. S2B, ESI†). These data demonstrate that bibacillin 1 has only weak activity against erythrocytes.

**Fig. 4.**
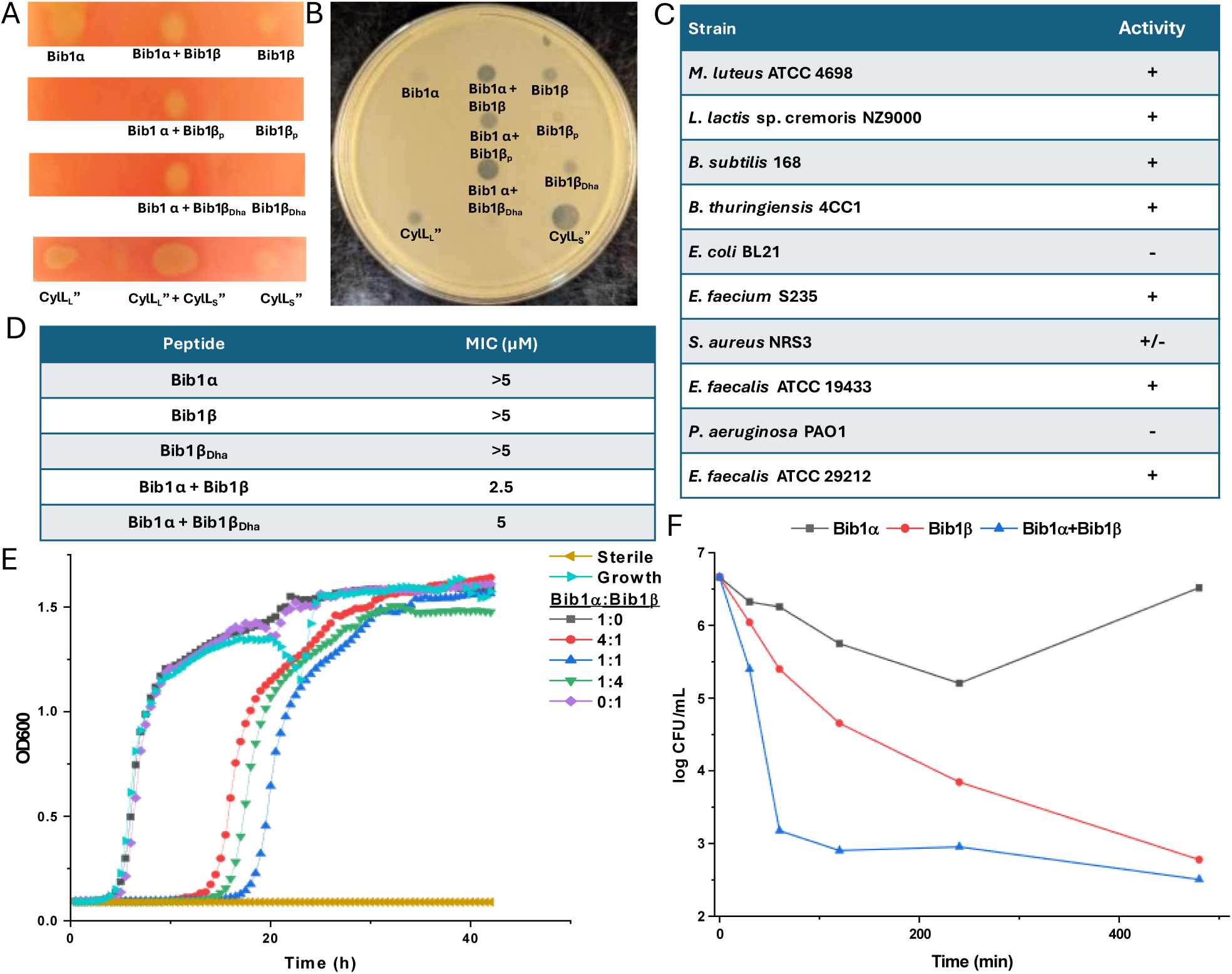
Activity of bibacillin 1 against erythrocytes and bacteria. A) Hemolytic activity of bibacillin 1 against rabbit erythrocytes. Cytolysin was used as a positive control. In each case, 300 pmol of peptide was spotted. B) Antibacterial activity of bibacillin 1 against *M. luteus*, with 300 pmol of peptide added to each indicated spot. C) Activity of bibacillin 1 (Bib1α:Bib1β 1:1) against at variety of Gram-positive bacteria including several ESKAPE pathogens. + indicates that a clear zone of growth inhibition was observed. – indicates that a zone of inhibition was not observed. +/- indicates that marginal inhibition was observed. D) Activity of Bib1α, Bib1β, Bib1β_Dha_, Bib1α + Bib1β (1:1) and Bib1α + Bib1β_Dha_ (1:1) in liquid growth assays. The minimum inhibitory concentration was defined as the lowest concentration of peptide at which sterility was observed 20 h after combining peptide with 5 x 10^5^ colony forming units (CFU) per mL of *M. luteus* culture. E) Growth of *M. luteus* treated with different compositions of Bib1α and Bib1β. Total peptide concentration was kept constant at 2.5 μM. F) Time-kill assay showing the change of CFU/mL of *M. luteus* cells over time after treatment with Bib1α, Bib1β or Bib1α:Bib1β (1:1). For each experiment, a total combined peptide concentration of 20 μM was used.

Using agar diffusion assays, we found bibacillin 1 to co-operatively inhibit the growth of *Micrococcus luteus* ATCC 4698 (Fig. 4B). Bib1α and Bib1β displayed weak individual activity in this assay whereas a stoichiometric mixture of these two peptides showed clear activity. Stoichiometric mixtures of Bib1α with Bib1β_P_ were less active than stoichiometric mixtures of Bib1α with Bib1β or Bib1β_Dha_. The structurally related peptide CylL_S_” showed strong activity that rivalled that of a 1:1 mixture of Bib1α:Bib1β. Stoichiometric mixtures of Bib1α and Bib1β showed micromolar activity under agar diffusion conditions against a wide range of Gram-positive bacteria, including members of the ESKAPE family of pathogens (Fig. 4C). Bibacillin 1 also displayed activity against *Enterococcus faecalis* ATCC 29212, a natural producer of cytolysin that possesses a gene encoding CylI, which is a protein that confers protection against cytolysin.^64^ This observation suggests that bibacillin 1 possesses an action mechanism that differs from that of cytolysin or that the recognition by CylI is sequence specific. In a broth dilution assay, Bib1α and β cooperatively inhibited the growth of *M. luteus* at concentrations as low as 2.5 µM when administered in a 1:1 ratio, and a stoichiometric mixture of Bib1α and Bib1β_Dha_ was similarly active with a minimal inhibitory concentration (MIC) of 5 µM (Fig. 4D). Under the same conditions, Bib1α, Bib1β and Bib1β_Dha_ did not show any individual activity at concentrations ≤5 μM. To compare the activity of bibacillin 1 to cytolysin, we also measured its MIC against *Lactococcus lactis* sp cremoris NZ9000, which is killed by cytolysin at concentrations as low as 32 nM.^35^ Bibacillin 1 exhibited an MIC of 5 μM against this strain. In combination, these data support a unique bioactivity profile for bibacillin 1 compared to cytolysin suggesting that bibacillin 1 has a different mechanism of action. The micromolar MICs observed for the individual peptides as well as the mixture are similar to the values reported previously for the single component cerecidins.^34^

We attempted to determine the ratio of Bib1α and Bib1β that gives maximal activity at a fixed total peptide concentration. As shown in Fig. 4E, a 1:1 ratio of Bib1α and Bib1β gives maximal activity at 2.5 μM but the activities at 1:4 and 4:1 were comparable. At higher concentrations (2xMIC), we observed maximal activity at 1:16, 1:8, 1:4 and 1:2 Bib1α:Bib1β (Fig. S3, ESI†). An agar diffusion assay with *M. luteus* showed that Bib1α:Bib1β ratios of 1:1, 1:2 and 1:3 produced nearly identical zones of growth inhibition (Fig. S3, ESI†). These results are not fully consistent with complexation of Bib1α and Bib1β, which should give a defined maximum in a continuous variation experiment. Instead, it appears that Bib1α and Bib1β may possess two different but complementary targets to which they bind with different affinities. Considering that higher activity seems to occur when Bib1β is in excess of Bib1α, it appears that only small amounts of Bib1α are necessary to fully sensitize bacteria to the effects of Bib1β. These observations are quite different from the activities reported for cytolysin, which displays a clear activity maximum at a 1:1 ratio,^65^ and all other currently characterized two-component lantibiotics.^24–26^

Time-kill assays were used to determine how rapidly the number of viable bacteria in a culture changed upon treatment with Bib1α, Bib1β, or a combination of both (Fig. 4F). Both Bib1α and Bib1β were bactericidal with Bib1β showing stronger activity. When Bib1α and Bib1β were combined in a 1:1 ratio, rapid cell death was observed, and minimal cell counts were documented 8 h after treatment demonstrating strong synergistic bactericidal activity. Bib1β reduced the cell counts to similar levels after 8 h but the killing occurred less rapidly. In contrast, cells began to recover from treatment with Bib1α after 4 h. The rapid killing induced by a stoichiometric mixture of Bib1α and Bib1β is consistent with cell lysis suggesting that bibacillin 1 may kill cells through co-operative pore formation.

The observed activity differences between Bib1α and Bib1β seem to be inconsistent with their structural similarities, but notions of similarity have previously failed to predict the activity of CylL_S_”-like sequences. For example, cerecidin is structurally very similar to CylL_S_” containing nearly the same number of residues, similar rings of the same size, and the same non-overlapping ring pattern; however, despite these similarities cerecidin is not co-operative with CylL_L_”.^34,35^ These results suggest that CylL_S_”-like peptides may have bioactivity profiles that are more diverse than indicated by their sequences. Considering that CylL_S_”-like sequences comprise one of the largest groups of class II lanthipeptides,^37^ these natural products may be a fruitful source of new antibiotics with diverse action mechanisms.

### Mechanism of Action Studies with Bibacillin 1

Two-component lantibiotics possess complex mechanisms of action that, to date, often defy complete explanation.^6^ The best understood two-component lantibiotics are lacticin 3147 and haloduracin, both of which possess a mersacidin-like component that binds to lipid II.^22,26,27^ Lipid II binding is thought to halt incorporation of the glycoside component of lipid II into the peptidoglycan layer, interrupting cell wall biosynthesis. This effect was shown in vitro with Halα, one of the components of haloduracin, which completely halted the consumption of lipid II by penicillin binding protein (PBP) when 2 equivalents of haloduracin relative to lipid II were present.^26^ In addition to blocking cell wall synthesis, haloduracin and lacticin 3147 show strong membrane depolarization activity, and studies involving model membranes doped with lipid II showed that lacticin 3147 forms pores more readily in membranes containing lipid II.^27^ Similarly, haloduracin rapidly depolarizes *B. subtilis* at concentrations that are coincident with its MIC.^25^

Although bibacillin 1 does not possess a mersacidin-like component, it does bear a resemblance to cacaoidin (Fig. S4, ESI†), which has a similar non-overlapping ring pattern and an N-terminal Dhx-Dhx-Xxx-Xxx-Cys sequence that is cyclized.^66^ Recently, cacaoidin was shown to target cell wall synthesis through a two-pronged attack involving lipid II binding and direct PBP inhibition.^67^ Based on the structural similarity to cacaoidin, we hypothesized that the bibacillin subunits might also target lipid II. To test this hypothesis, we recombinantly expressed and purified *E. coli* PBP1b following a previously described procedure,^68^ yielding active PBP1b that readily consumed biosynthesized samples of Gram-positive lipid II (Fig. 5A). Consistent with previous observations, a thin layer chromatography experiment demonstrated that four equivalents of Halα protected lipid II from consumption by PBP1b. Under the same conditions, Bib1α and Bib1β were incapable of protecting lipid II (Fig. 5A). When Bib1α and Bib1β were administered together in a 1:1 ratio, lipid II was also not protected from consumption by PBP1b. These results suggest that bibacillin 1 does not target lipid II in a manner that blocks cell wall synthesis, distinguishing it from other investigated two-component lantibiotics.

**Fig. 5.**
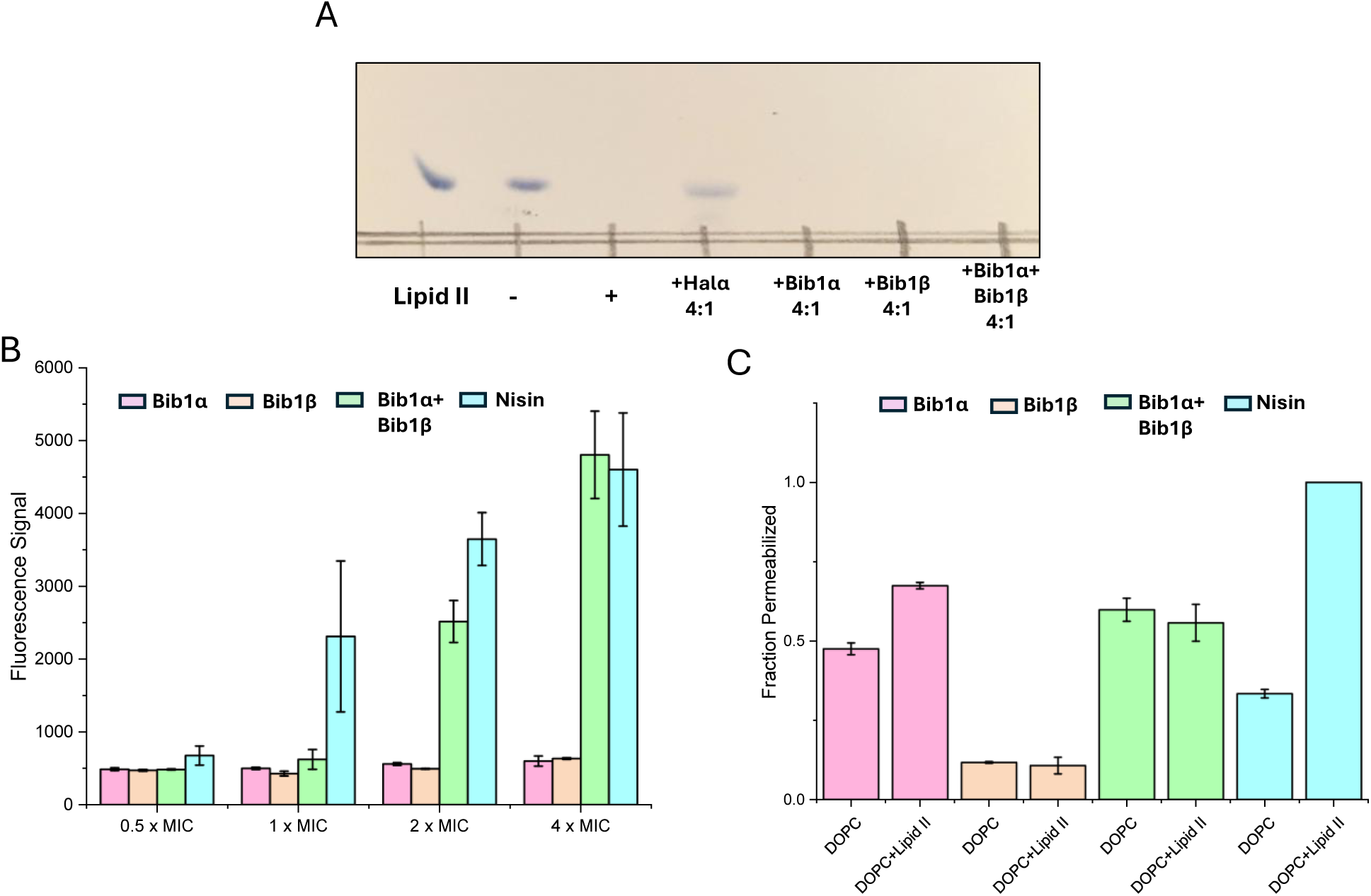
Bibacillin 1 does not target lipid II and does not cause rapid membrane depolarization at its MIC. A) Thin layer chromatography showing consumption of lipid II in a PBP1b reaction and the effects of the presence of 4 equivalents of peptide relative to lipid II. The lipid II lane contained 0.1 nmol of purified lipid II starting material. +/- indicates the presence/absence of PBP1b. The plate was stained with cerium ammonium molybdate after development. B) DiSC_3_(5) fluorescence signal after incubation of *M. luteus* (OD_600_ = 0.2) with Bib1α or Bib1β individually or as a 1:1 mixture for 30 min. For each concentration indicated, the total amount of bibacillin peptide was kept constant (MIC = 2.5 μM). Nisin was used as a positive control and its concentration was scaled in accordance with its MIC (1 μM). C) Pore formation activity of Bib1α, Bib1β, a stoichiometric mixture of Bib1α and Bib1β, and nisin in the presence of liposomes comprised of DOPC containing or lacking 0.1 mol% lipid II. In all cases, a total peptide concentration of 5 μM was used.

As mentioned, two-component lantibiotics often display pore formation activity and this pore formation activity can be lipid II dependent. To determine whether bibacillin 1 is capable of depolarization of *M. luteus* cells at its MIC, we used the membrane potential sensitive dye DiSC_3_(5).^69^ Using nisin as a positive control, the membrane potentials of cells incubated with Bib1α or Bib1β individually or as a 1:1 ratio were monitored over the course of 30 min. Individually, Bib1α or Bib1β did not show strong membrane depolarization activity at any of concentrations tested. When Bib1α and Bib1β were combined in a 1:1 ratio, membrane depolarization activity similar to nisin was observed at concentrations 2- to 4-fold higher than the MIC, but very little activity was observed at the MIC. In contrast, nisin showed half-maximal pore formation activity at its MIC. These results suggest that bibacillin 1 can engage in cooperative membrane depolarization, but this activity appears not relevant at its MIC.

To investigate the lipid II-dependence of the pore formation activity of bibacillin 1, pyranine-encapsulated liposomes were prepared containing or lacking lipid II and pore formation was monitored via pH-dependent pyranine fluorescence.^70^ Consistent with the literature, the pore formation activity of nisin was strongly dependent on the presence of lipid II in the assayed membranes (Fig. 5B).^71^ At 5 μM, Bib1α showed pore formation activity that was somewhat enhanced by the addition of lipid II. In contrast, Bib1β did not show pore formation activity at 5 μM. When Bib1α and Bib1β were combined in a 1:1 ratio they showed pore formation activity similar to Bib1α but the presence of lipid II did not enhance pore formation activity. These results are inconsistent with bibacillin 1 participating in lipid II-dependent pore formation.

Experiments with bacterial cells described above demonstrated that the membrane depolarization of bibacillin 1 was cooperative at high concentrations (Fig. 5B), which is in contradiction with the pore formation studies conducted using liposomes where no cooperativity or slight anti-cooperativity was observed at the same concentrations (Fig. 5C). These results suggest that bibacillin 1 does not form pores at its MIC and likely kills cell through direct targeting of a certain cellular process; however, at higher concentrations (≥2 x MIC), complexes of bibacillin 1 with its target may be able to assemble into oligomeric pores resulting in rapid cell death. This activity is reminiscent of daptomycin, which kills cells through binding to phosphatidylglycerol in a manner that inhibits cell wall synthesis, that also shows pore formation activity 2-4 fold above its MIC, but not at its MIC.^72–74^

### Quorum sensing activity of bibacillin 1

The quorum sensing system found in the cytolysin BGC is fascinating and enigmatic. It was previously shown that CylL_S_” could stimulate the production of cytolysin through an interaction with CylR1, a small protein thought to be embedded in the cellular membrane, which influences the DNA binding of the transcription factor CylR2.^75^ Unlike other quorum sensing systems found in *E. faecalis*, signal transduction does not proceed through a phosphorylation relay.^76^ The cytolysin quorum sensing system ties cytolysin expression to local cellular density through an aggregation mechanism. It is thought that CylL_S_” and CylL_L_” undergo aggregation in the absence of proximal cell membranes.^43^ When the cell density reaches a certain level, CylL_L_” becomes embedded into the encroaching cell membranes, liberating CylL_S_” for stimulation of cytolysin expression. To date, the cytolysin quorum sensing system has only been described for the enterococcal cytolysin and the prevalence of this quorum sensing system in other BGCs has not been explored. The CylR1/R2 analogs found in the bibacillin 1 and bibacillin II BGCs are almost identical to each other, and the single component CylL_S_”-like lantibiotic cerecidin does not possess a CylR1/R2 system, suggesting that the system may be specific for two-component lantibiotics. However, carnolysin,^77^ a two-component class II lantibiotic from *Carnobacterium maltaromaticum* that is structurally similar to cytolysin, does not possess a CylR1/CylR2-like system, suggesting that this quorum sensing system is not widely distributed in the BGCs of cytolysin-like two-component lantibiotics.

Considering the structural similarities between CylL_S_”, Bib1α and Bib1β, and the presence of genes encoding BibR1 and BibR2 in the BGC, we speculated that bibacillin 1 could be involved in a similar quorum sensing mechanism. Furthermore, these peptides may also conduct interspecies crosstalk or influence the fitness of neighbouring cells that contain similar quorum sensing systems. To assess the latter hypothesis, we tested the ability of bibacillin 1 to trigger the CylR1/R2 system using a previously described reporter strain *E. faecalis* pXL110.^40^ This strain contains a lacZ gene under CylR1/R2 control; thus, CylR1/R2 triggering can be inferred from LacZ activity.

Using agar plates seeded with *E. faecalis* pXL110 and the colorimetric LacZ substrate X-gal, LacZ expression was visualized. Consistent with previous reports,^43^ CylL_S_” stimulated LacZ expression when 30 pmol of the peptide was deposited onto the agar plate. In contrast, nisin did not induce LacZ expression even when applied at a 10-fold higher concentration compared to CylL_S_”. These results demonstrated that the assay is specific for CylL_S_”-like peptides and cell killing by nisin does not result in non-specific triggering of CylR1/R2 (Fig. 6A). Bib1α, Bib1β and Bib1β_P_ (Fig. S5, ESI†) did not stimulate LacZ expression. Comparing the structures of CylL_S_” and Bib1β (Fig. 1A and 2B), we speculated that the presence of Ser16 in Bib1β rather than the hydrophobic residues present in CylL_S_” in this region might interfere with CylR1 binding possibly due to hydrophobic mismatching. To investigate this hypothesis, we tested Bib1β_Dha_ (Fig. 6A) and Bib1β S16A (Fig. S5, ESI†) and found that only Bib1β_Dha_ stimulated the CylR1/R2 system. Interestingly, when Bib1β_Dha_ was combined in a 1:1 ratio with Bib1α LacZ expression was no longer observed suggesting that, like cytolysin, bibacillin 1 may cooperatively aggregate in a manner that prevents Bib1β_Dha_ from stimulating BibR1/R2. To determine whether bibacillin 1 could quorum sense through an aggregation mechanism, the aggregation propensity of bibacillin 1 was analyzed using SDS-PAGE and dynamic light scattering (DLS). Previously, stoichiometric mixtures of CylL_S_” and CylL_L_” were found to assemble into oligomers with masses of 130-220 kDa and this oligomerization event was implicated in the reversible sequestration of CylL_S_” that is thought to prevent CylR1/R2 stimulation.^43^ Performing a similar experiment on bibacillin 1, SDS-PAGE analysis of Bib1α and Bib1β revealed that they are oligomeric in the presence of SDS, but this oligomerization event is not co-operative (Fig. S6, ESI†). Moreover, unlike the observations with cytolysin,^43^ oligomers in the 130-220 kDa range were not observed suggesting that sequestration of Bib1β_Dha_ occurs via a different mechanism. Since SDS can influence aggregation, the aggregation of bibacillin 1 was further investigated under SDS-free conditions using DLS (Fig. 6B). Bib1α exists in solution as aggregates that are 159 nm in diameter on average (PDI = 0.224) and Bib1β forms aggregates that are 291 nm in diameter on average (PDI = 0.384). The size distribution of the Bib1β aggregates is bimodal suggesting the presence of multiple oligomeric forms. When the two subunits were mixed in a 1:1 ratio, aggregates of intermediate sizes (diameter = 182 nm; PDI = 0.254) were detected suggesting that the aggregation state may not be co-operative. Taken together, these data suggest that bibacillin 1 quorum sensing involves a mechanism that does not depend on cooperative aggregation. In-lieu of a cytolysin-like aggregation mechanism, it is possible that Bib1α and Bib1β function as an antagonist-agonist pair that compete for the BibR1/BibR2 system receptor effectively tying the expression of bibacillin 1 to the ratio of these two peptides. Confirmation of this hypothesis will require in vitro studies with Bib1R1/R2.

**Fig. 6.**
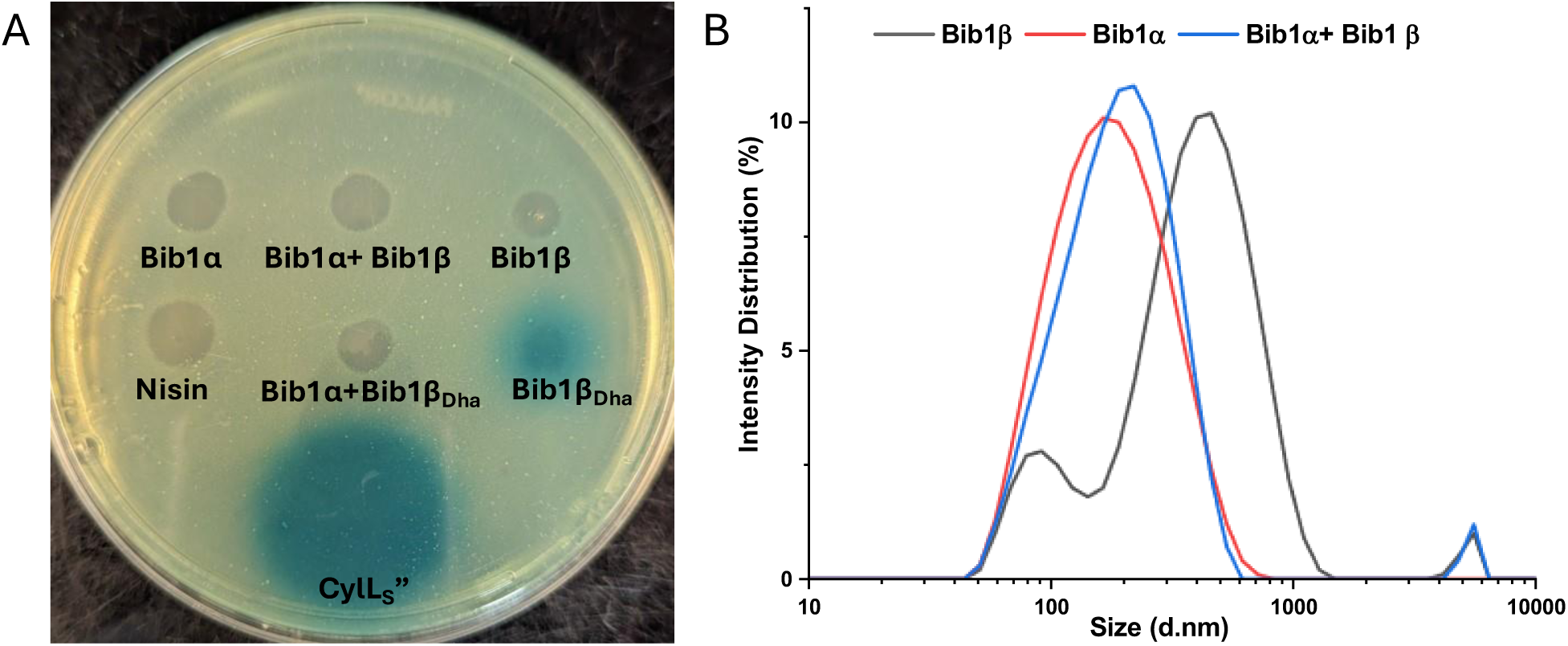
Exploring the quorum sensing mechanism of bibacillin 1. A) Assays to assess whether bibacillin 1 triggers the CylR1/R2 system. A reporter strain of *E. faecalis* with LacZ expression under the control of the CylR1/R2 system was seeded into soft agar containing X-gal. Next, 300 pmol of Bib1α, Bib1β, and Bib1β_Dha_ either separately or as 1:1 mixtures were deposited onto the soft agar layer. Nisin (300 pmol) and CylL_S_” (30 pmol) were used as negative and positive controls, respectively. B) Size distributions obtained using dynamic light scattering showing the oligomeric states of Bib1α, Bib1β, and Bib1α:Bib1β (1:1) in phosphate buffered saline (pH = 7.4) at room temperature. In each case, the total peptide concentration was 1 μM.

Stimulation of the CylR1/R2 system by a non-cognate peptide suggests that CylL_S_”-like peptides could impact the fitness of neighbouring bacteria that possess CylR1/R2-like systems through either over stimulating the system, resulting in unnecessary peptide production, or antagonising the system, decreasing fitness at high cellular densities. This observation also suggests that co-colonization of the gut microbiome with organisms that produce CylL_S_”-like peptides could impact cytolysin expression in a manner that impacts cytolysin-induced disease states.^33^ The extent to which this phenomenon impacts microbiomes requires further investigation.

The observed bioactivity of Bib1β, Bib1β_P_, and Bib1β_Dha_ leaves some uncertainty regarding the identity of the truly mature form of modified Bib1A2. Bib1β_P_ showed reduced antibacterial activity compared to Bib1β and Bib1β_Dha_, and it did not stimulate CylR1/R2 suggesting that it was an intermediate form of the true natural product. However, in light of previous reports that have shown that class II lanthipeptide synthetases produce the same product in *E. coli* as in the native producer, the production of significant quantities of phosphorylated Bib1A2 may have a physiological role as demonstrated for bacteria in the oral microbiome.^11^ The antibacterial activities of Bib1β_Dha_ and Bib1β were nearly identical when they were combined with Bib1α, but Bib1β_Dha_ stimulated CylR1/R2 whereas Bib1β did not. Taken together, it is tempting to conclude that the mature form is Bib1β_Dha_; however, although BibR1 and CylR1 have similar sequences (30% identical and 51% similar), in the absence of a detailed characterization of the CylL_S_”-CylR1 interaction it is difficult to conclude whether recognition by one implies recognition by the other.

## Conclusions

Bibacillin 1 is a new entry into the growing collection of two-component lantibiotics that are structurally different from the prototype lacticin 3147.^29,30,57^ Co-expression of Bib1A1 and Bib1A2 with their cognate synthetase in *E. coli* yielded modified peptides that were converted to their mature counterparts through treatment with the protease CylA. The mature peptides have a non-overlapping ring pattern and are more similar to CylL_S_” than CylL_L_”. They both possess a LL-MeLan A ring and a DL-Lan B ring, consistent with the presence of a Dhx-Dhx-Xxx-Xxx-Cys at the N-terminus of the Bib1α/β sequence, which was previously shown to favour the formation of LL-MeLan. During the co-expression of Bib1A2 with Bib1M, significant fractions of Bib1A2 were phosphorylated at Ser64 (Ser16 in Bib1β) leaving some ambiguity about the true identity of the mature form of Bib1A2. To address this, we also isolated the phosphorylated form of Bib1β and devised a route to the fully dehydrated Bib1β_Dha_.

Bibacillin 1 lacked the strong hemolytic activity possessed by the enterococcal cytolysin, displayed co-operative antibacterial activity against a collection of Gram-positive organisms, and was inactive against Gram-negative organisms. Bib1β_P_ displayed lower antibacterial activity compared to Bib1β and Bib1β_Dha_ when combined with Bib1α suggesting that Bib1β_P_ was an intermediate during the formation of Bib1β_Dha_ or serves some other purpose to the natural producer.^11^ The impact of bibacillin 1 on the growth of *M. luteus* under continuous variation conditions demonstrated that, unlike other examples of two-component lantibiotics, Bib1α and Bib1β are unlikely to form a discrete complex as part of their action mechanism. These results are consistent with a model in which each peptide may bind to a different target with Bib1α serving to sensitize the bacteria to attack by Bib1β. Time-kill assays showed that both peptides are individually bactericidal while a stoichiometric mixture of Bib1α and Bib1β caused cell death that was significantly more rapid.

Unlike lacticin 3147, haloduracin, and cacaoidin, bibacillin 1 does not kill cells through a lipid-II dependent mechanism. The results from pore formation studies are also inconsistent with bibacillin 1 killing cells through cooperative pore formation at its MIC, although the peptides did engage in cooperative membrane depolarization at concentrations at least 2-fold higher than the MIC.

The *bib* BGC encodes two proteins with similarity to CylR1/R2 suggesting that the peptides may be involved in quorum sensing and could serve to impact natural product production in neighbouring bacteria. Only Bib1β_Dha_ stimulated the CylR1/R2 system and combining Bib1β_Dha_ with Bib1α in a 1:1 ratio eliminated this ability suggesting that the sensor system could be affected by co-operative peptide aggregation. However, investigations into the aggregation of bibacillin 1 did not show evidence of cytolysin-like oligomerization and aggregation events suggesting that bibacillin 1 may engage in a different quorum sensing mechanism. The enhanced bioactivity of Bib1β_Dha_ compared to Bib1β and Bib1β_P_ suggests that Bib1β_Dha_ is likely the mature form of the natural product with Bib1β_P_ serving as an intermediate form. This assignment is predicated on the assumption that CylR1/R2 functions as a good model for Bib1R1/R2 and the strength of this assumption is difficult to ascertain currently.

Overall, bibacillin 1 furthers a growing vein of literature which shows that CylL_S_”-like sequences display more diverse bioactivities than would be expected based on their sequence similarities.

## Experimental

### General Methods

Enzyme mixtures used for molecular biology were purchased from New England BioLabs (NEB) (Ipswich, MA). Chemicals were purchased from Sigma-Aldrich (St. Louis, MO). Growth media were purchased from Becton, Dickinson and Company (Franklin Lake, NJ) or Fisher Scientific (Pittsburgh, PA). Single stranded DNA used as primers for PCR were purchased from Integrated DNA Technologies (Newark, NJ). Gene fragments were purchased from Twist Bioscience (San Francisco, CA). The sequences of primers and gene fragments can be found in Table S2 and Table S3, respectively. Whole plasmid sequencing was performed by Plasmidsaurus using Oxford Nanopore Technology with custom analysis and annotation. Matrix-assisted laser desorption/ionization time-of-flight mass spectrometry (MALDI-TOF MS) analysis was performed on a Bruker UltrafleXtreme MALDI-TOF mass spectrometer (Bruker Daltonics) at the University of Illinois School of Chemical Sciences Mass Spectrometry. All samples analyzed by MALDI-TOF MS were co-crystallized with Super DHB MALDI matrix purchased from Sigma-Aldrich. Semi-preparative reversed phase HPLC (R-HPLC) was performed on an Agilent 1260 infinity II LC system. Fluorescence emission intensities and optical densities were determined using a Biotek Synergy H4. Dynamic light scattering (DLS) analysis was performed on a Zetasizer Nanoseries (Malvern Panalytical,). CylL_S_” and CylL_L_” were prepared using a previously described procedure.^35^ Nisin was isolated from commercially available crude extracts of *Lactococcus lactis* (Sigma Aldrich).

### Strains and Plasmids

*E. coli* DH5α was used as host for cloning and plasmid propagation. *E. coli* BL21 (DE3) served as a host for overexpression of proteins and peptides. The pRSFDuet-1 co-expression vector was obtained from Novagen. Strains used in bioactivity and mechanism of action studies are listed in Table S1 along with their growth conditions.

### Bibacillin 1 Expression Constructs

Gene fragments containing the codon optimized nucleotide sequence of N-terminal His-tagged *bib1A1* or *bib1A2* were purchased from Twist Biosciences (Table S3) and amplified by PCR using primers with homologous sequences to multiple cloning site 1 in pRSFDuet-1 (Bib1A1 FP and Bib1A1 RP; Bib1A2 FP and Bib1A2 RP. See Table S2). Two codon optimized gene fragments corresponding to two portions of *bib1M* were purchased from Twist Biosciences (Table S3) and amplified by PCR using one primer with a homologous sequence to multiple cloning site 2 in pRSFDuet-1 (Bib1A1Mfrag1 FP and Bib1A1Mfrag2 RP or Bib1A2Mfrag1 FP and Bib1A2Mfrag2 RP. Table S2) and another primer with a homologous sequence to the other fragment (Bib1A1Mfrag1 RP and Bib1A1Mfrag2 FP or Bib1A2Mfrag2 RP and Bib1A2Mfrag2 FP. Table S2). pRSFDuet-1 was linearized through amplification by PCR using primers flanking the insertion sites (Bib1A1 pRSF RP and Bib1A1 pRSF FP or Bib1A2 pRSF FP and Bib1A2 pRSF RP. Table S2). Amplified fragments were isolated via agarose gel purification using a NucleoSpin Gel and PCR Clean-up kit (Macherey-Nagel). The fragments were then combined using Gibson Assembly^®^ Master Mix (NEB) and cloned. Co-expression constructs for the expression of Bib1A2-S64C or Bib1A2-S64A along with Bib1M were produced through single site mutagenesis of the His_6_-Bib1A2:Bib1M:pRSF-Duet1 plasmid. End-to-end primers were designed with the mutation present near the middle of one of the primers (Bib1A2 SSM S64C FP and Bib1A2 SSM S64C RP or Bib1A2 SSM S64A FP and Bib1A2 SSM S64A RP). Whole plasmid amplification by PCR followed by treatment with KLD Enzyme Mix (NEB) yielded a sample of circularized plasmid that was directly used for cloning. Plasmid sequences were verified by whole plasmid sequencing.

### Overexpression and Purification of Peptide

Overnight cultures of *E. coli* BL21 (DE3) containing the plasmid of interest were prepared in Lysogeny Broth (Lennox) containing 50 μg/mL kanamycin and diluted 50:1 into Terrific Broth (TB) containing 50 μg/mL kanamycin. The cultures were grown at 37 °C until an OD_600_ of 0.8 was reached. The cultures were then cooled in ice-water for 30 min before induction with IPTG (0.3 mM), followed by expression for 13 h at 16 °C. Cells were then harvested by centrifugation (6000 x g for 12 min at 4 °C) and resuspended in lysis buffer (6.0 M guanidinium hydrochloride, 0.5 mM imidazole, 0.5 M NaCl, 20 mM NaH_2_PO_4_, pH 7.5) at a concentration of 5 mL of lysis buffer for every 1 g of cell pellet. The suspended cells were placed on a rotator and gently mixed for 30 min at 4 °C followed by sonication. The lysate was centrifuged for 40 min at 12000 g at 4 °C before incubation with His60 SuperFlow resin (Takara Bio) for 1 h at 4 C°. Approximately 1 mL of resin was used for every 15 g of cell pellet. The resin was collected by centrifugation (1000 g for 5 min at 4 °C) and the supernatant was discard. The resin was transferred to a fritted vessel and washed with 3 bed volumes (BV) of lysis buffer. The resin was further washed with 10 BV each of Lan A buffer B2 (4.0 M guanidinium hydrochloride, 30 mM imidazole, 0.3 M NaCl, 20 mM NaH_2_PO_4_, pH 7.5) and LanA co-expression wash buffer (30 mM imidazole, 0.3 M NaCl, 20 mM NaH_2_PO_4_, pH 7.5). Peptide bound to the resin was eluted with 10 BV of elution buffer (0.5 M imidazole, 0.3 M NaCl, 20 mM NaH_2_PO_4_, pH 7.5).

### Peptide Cleavage and HPLC Purification

Peptide in elution buffer was treated with freshly prepared CylA (40 nM) for 6-10 h at room temperature. The completeness of the cleavage reaction was checked by desalting a small sample using Peptide Cleanup C18 Pipette Tips (Agilent) and analysis using MALD-TOF MS. Once complete, the peptide sample was acidified with trifluoroacetic acid (TFA) to a pH of 4. The acidified samples were clarified by centrifugation (4000 g for 10 min) then purified by R-HPLC using a Phenomenex Jupiter Proteo column (4 μm, 90 Å, 10 × 250 mm) column. A gradient method was used for peptide purification which employed mixtures of H_2_O + 0.1% TFA (solvent A) and CH_3_CN + 0.1% TFA (solvent B) and a flow rate of 4 mL/min. The gradient started with 2% B for 10 min then proceeded to a linear ramp of solvent B to 85% over 25 min. Product was detected using MALDI-TOF MS and product containing fractions were combined and lyophilized. Product purity was assessed via LCMS-MS.

### Conversion of His_6_-Bib1A2 S64C to Bib1β_Dha_

Modified His_6_-Bib1A2-S64C was prepared by co-expression with Bib1M following the overexpression and purification procedure. Once eluted from the nickel column, the peptide was desalted using a Bond Elut C18 solid phase extraction cartridge and eluted with 60% CH_3_CN + 0.1% TFA and H_2_O + 0.1% TFA. Peptide containing fractions were determined using MALDI-TOF MS then combined and lyophilized. The lyophilized peptide was dissolved in water and diluted to a concentration of 2 mM. Tris(2-carboxyethyl) phosphine (TCEP) was then added to a concentration of 0.5 mM and the solution was incubated at room temperature for 30 min before diluting with an equal volume of DMSO containing 2,5-dibromomethylvalerate (60 equiv.). The mixture was immediately transferred to a separate vessel containing solid K_2_CO_3_ (150 equiv.) and incubated at 37 °C for 4 h before reaction completeness was assessed using MALDI-TOF MS. Once complete, the solution was diluted 12-fold with phosphate buffered saline (PBS; pH = 7.4) and an Amicon with a 3 kDa molecular weight cutoff was used to exchange the peptide into PBS. The peptide was then treated with CylA (40 nM) at room temperature for 6-10 h. The solution was acidified with TFA to pH = 7.4 and purified using the R-HPLC conditions described in the peptide cleavage and HPLC purification section.

### High-Resolution Tandem Mass Spectrometry

Purified samples of peptide were injected into an Agilent 1290 LC-MS QToF instrument for HR-MS/MS analysis. LC separation occurred at a temperature of 45 °C using a 10 min gradient proceeding linearly from 5% CH_3_CN+0.1% formic acid (FA)/95% H_2_O+0.1% FA to 95% CH_3_CN+0.1% FA/5% H_2_O+0.1% FA and a flow rate of 0.4 mL/min was used throughout. For all peptides analyzed in this study, a Kinetex® C8 (2.6 μm, 100 Å 2.1 x 150 mm) LC column was used. Tandem-MS fragmentation employed normalized collision energies of 20 and 30.

### Agar Diffusion Hemolysis Assay

Defibrinated rabbit erythrocytes (Hemostat) were washed by 5-fold dilution in cold sterile PBS, followed by centrifugation at 1000 g for 10 min and decanting. This process was repeated until the supernatant was clear. The washed erythrocytes were diluted to a concentration of 5% (v/v) in molten BHI containing 0.7% agar cooled to 50 °C and the mixture was immediately deposited onto a warm 2% BHI agar plate. After setting at room temperature for at least 30 min, 3 μL of peptide solution diluted in PBS to a concentration of 100 μM was deposited onto the agar plate and the plate was allowed to stand for another 30 min. The plate was then incubated at 37 °C for 16 h before documentation.

### Solution Phase Hemolysis Assay

Solution phase hemolysis assays were conducted as described previously.^35^

### Agar Diffusion Assays using Bacteria

Overnight cultures of bacteria were prepared according to Table S1. The OD_600_ of each culture was determined and adjusted to a value of 1 using the appropriate growth media. Molten growth media containing 0.7% agar was cooled to 50 °C and inoculated with 50 μL of adjusted overnight culture for every 5 mL of molten growth media. The inoculated molten growth media was immediately poured onto a pre-warmed agar (2%) plate prepared using the same media. The plate was allowed to set for 30 min at room temperature before adding 3 μL of peptide stock diluted in PBS to 100 μM. The deposited peptide solutions were allowed to stand on the agar plate for 30 min before incubation for 16-36 h at the optimum growth temperature until the desired cell density was achieved.

### Time-Kill Assay

An overnight culture of *M. luteus* ATCC 4698 was adjusted to approximately 10^6^ CFU/mL and 20 μM of Bib1α, Bib1β or Bib1α:Bib1β was added. The solutions were incubated at 30 °C with shaking. At each time point, 20 μL was removed from each culture and immediately streaked onto tryptic soy broth agar plates. The plates were incubated at 30 °C for 36-48 h and colonies were counted.

### Continuous Variation Growth Curve

Mixtures of Bib1α and Bib1β in growth media were prepared on a sterile 96-well plate keeping the total peptide concentration constant in each row of the plate. The samples in each well were serially diluted (2-fold) and then an equal volume of *M. luteus* ATCC 4698 suspended in growth media was added to 5 x 10^5^ CFU/mL. The plate was transferred to a plate reader preheated to 30 °C and shaken continuously. The OD_600_ of each well was measured every 30 min for 48 h.

### In Vitro Peptidoglycan Synthesis Assay with Lipid II Substrate

A solution of lipid II (1.5 nmol) in 1:1 CHCl_3_:MeOH that was prepared by a previously disclosed procedure^78^ was added to a low binding Eppendorf tube and concentrated to remove any organic solvent. Triton X-100 (1% v/v, 2 uL) was added to the residue and the mixture was sonicated briefly before diluting with 37.5 μL of reaction buffer (20 mM MES, 2 mM MgCl_2_, 2 mM CaCl_2_, pH = 5.5). The solution was then transferred to a new low binding Eppendorf tube containing lanthipeptide. The mixture was vortexed and briefly sonicated to ensure dissolution. The mixtures were incubated for 10 min at room temperature before adding PBP1b (6 μg), which was prepared as described elsewhere,^68^ and then incubated at 30 ° C. After 2 h, 150 μL nBuOH:AcOH-pyridine (2:1) was added to the reaction mixture and the organic layer was removed, washed with 40 μL of water and then concentrated. The residue was dissolved in 12 µL of 1:1 MeOH:CHCl_3_ and 3 μL was deposited onto a silica gel plate. The TLC plate was developed in mobile phase (88:48:10:1 CHCl_3_:MeOH:H_2_O:NH_4_OH) for 10 min before drying and staining the plate with cerium ammonium sulfate. The plate was documented immediately after the staining procedure was completed.

### Membrane Depolarization Assay

An overnight culture of *M. luteus* ATCC 4698 was diluted to an OD_600_ of 0.2 and bovine serum albumin (1 mg/mL) and DiSC_3_(5) (1 μM) were added. The mixture was transferred to a 96-well black plate and the fluorescence of DiSC_3_(5) was measured using an excitation wavelength of 622 nm and an emission wavelength of 670 nm throughout the experiment. The fluorescence signal was measured for 8 min prior to the addition of peptides. After adding peptides, the fluorescence signal was monitored for 30 min. The maximum fluorescence signal achieved over the entire course of the experiment was considered to be a metric of membrane depolarization activity and was plotted as a function of peptide concentration.

### In Vitro Pore Formation Assay

This assay was conducted as described elsewhere.^70^

### CylR1/R2 Triggering

Colorimetric agar diffusion assays were used to determine CylR1/R2 triggering. Briefly, 0.7% BHI soft agar was mixed with X-gal (1.6 mg/mL of molten agar) seeded with an overnight culture of the *Enterococcus faecalis* FA2-2 (pLX110) reporter strain (200 µL of overnight culture for every 5 mL of molten agar). After setting for 30 min at room temperature, peptides were added to the soft agar layer and incubated at room temperature for 30 min. The plates were then incubated overnight at 37 °C and documented.

## Author Contributions

R.M. and W.A.v.d.D conceptualized this study and wrote the manuscript. R.M. designed and executed the experiments. Y.Y. assisted with the preparation and purification of materials. Y.L. performed Marfey’s analysis. M.S.G. was a source for insightful discussion and provided the reporter strain pLX110. All authors edited the manuscript.

## Conflicts of Interest

There are no conflicts to declare.

## Acknowledgements

This study was supported by the National Institutes of Health (R37GM058822 to W.A.v.d.D. and AI083214 to M.S.G.). The Bruker UltrafleXtreme MALDI TOF/TOF mass spectrometer was purchased in part with a grant from the National Center for Research Resources, National Institutes of Health (S10 RR027109 A). W.A.v.d.D. is an Investigator of the Howard Hughes Medical Institute (HHMI). R.M. was supported by a postdoctoral fellowship from the Life Sciences Research Foundation and a postdoctoral fellowship from the Natural Sciences and Engineering Research Councill of Canada. Y.Y was supported by a Spudich Undergraduate Research Scholarship in Chemistry. We thank Dr. Chandrashekhar Padhi for his help with the molecular biology featured in this work. This study is subject to HHMI’s Open Access to Publications policy. HHMI laboratory heads have previously granted a nonexclusive CC BY 4.0 license to the public and a sublicensable license to HHMI in their research articles. Pursuant to those licenses, the author-accepted manuscript of this article can be made freely available under a CC BY 4.0 license immediately upon publication.

## Notes

### Competing Interest Statement

The authors have declared no competing interest.

