## Supplementary Information for "Bibacillin 1: A two-component lantibiotic from *Bacillus thuringiensis*"


| | |
|---|---|
| Table S1. Strain information and growth conditions | S2 |
| Table S2. List of primers | S3 |
| Table S3. Sequences of codon optimized gene fragments | S3-4 |
| Table S4. Accession numbers | S5 |
| Fig. S1. LCMS-MS analysis of Bib1$\beta_P$ | S6 |
| Fig. S2. Solution phase hemolytic activity of bibacillin 1 | S7 |
| Fig. S3. Additional continuous variation experiments | S8 |
| Fig. S4. Structure of cacaoidin | S8 |
| Fig. S5. Bib1$\beta_P$ and Bib1β S16A CylR1/R2 triggering | S9 |
| Fig. S6. SDS-PAGE analysis of bibacillin 1 | S9 |
| Fig. S7. R-HPLC traces of purified Bib1α, Bib1β, Bib1$\beta_P$, Bib1$\beta_{Dha}$, and Bib1β S16A | S10 |
| Fig. S8. HRMS spectra of purified Bib1α, Bib1β, Bib1$\beta_P$, Bib1$\beta_{Dha}$, and Bib1β S16A | S11 |
| Table S5. Comparing high resolution mass to expected masses for Bib1α, Bib1β, Bib1$\beta_P$, Bib1$\beta_{Dha}$, and Bib1β S16A | S12 |
| References | S12 |

Table S1. Strain information and growth conditions

| Strain | Source | Media | Temperature (℃) |
| --- | --- | --- | --- |
| *Lactococcus lactis* subsp. cremoris NZ9000 | Kuipers et al.[1] | M17+0.5% glucose | 30 |
| *Micrococcus luteus* | ATCC[a] 4698 | Tryptic Soy Broth (TSB) | 30 |
| *Bacillus subtilis* 168 | Helmann et al.[2] | Lysogeny broth (Miller) | 37 |
| *Bacillus thuringiensis* subsp. pulsiensis | BGSC[b] 4CC1 | 3 nutrient broth (3N) | 30 |
| *Escherichia coli* BL21 (DE3) | UIUC-CMF[c] | Lysogeny broth or Terrific broth | 37 |
| *Enterococcus faecium* | ATCC 19434 | BHI | 37 |
| *Enterococcus faecalis* | ATCC 19433 | BHI | 37 |
| *Enterococcus faecalis* | ATCC 29212 | BHI | 37 |
| *Pseudomonas aeruginosa* PA0162 | Manoil et al.[3] | 3N | 37 |
| *Staphylococcus aureus* NRS3 | NARSA[d] | TSB | 37 |

[a]ATCC: American type culture collection. [b]BGSC: Bacillus genetic stock center. [c]UIUC-CMF: University of Illinois at Urbana-Champaign cell and media facility. [d]NARSA: Network on antimicrobial resistance in *Staphylococcus aureus*.

Table S2. Primers used for molecular biology.

| **Name** | **Sequence** |
|---|---|
| Bib1A1 FP | gtttaactttaataaggagatataccatgccacatcaccatca |
| Bib1A1 RP | tttgattcatggtatatctccttgaatcttagcagtgcatccggct |
| Bib1A1Mfrag1 FP | actgctaagattcaaggagatataccatgaatcaaaatacaagcataaaac |
| Bib1A1Mfrag1 RP | ttgccgttcagctttggcacgttgttatcgtttatatcagccataactt |
| Bib1A1Mfrag2 FP | atggctgatataaacgataacaacgtgccaaagctgaacggcaa |
| Bib1A1Mfrag2 RP | cggtttctttaccagactcgagctaatatttgataggcttttctaaggtcaa |
| Bib1A1 pRSF FP | cttagaaaagcctatcaaatattagctcgagtctggtaaagaaaccg |
| Bib1A1 pRSF RP | tgatggtgatgtggcatggtatatctccttattaaagttaaac |
| Bib1A2 FP | gtttaactttaataaggagatataccatgccgcatcatcac |
| Bib1A2 RP | ttatgcttgtattttgattcatggtatatctccttgaatctcaacaataaattttggagat |
| Bib1A2Mfrag1 FP | tttattgttgagattcaaggagatataccatgaatcaaaatacaagcataaaacccc |
| Bib1A2Mfrag1 RP | ttgccgttcagctttggcacgttgttatcgtttatatcagccataactttttttgcg |
| Bib1A2Mfrag2 FP | atggctgatataaacgataacaacgtgccaaagctgaacggcaa |
| Bib1A2Mfrag2 RP | ttctttaccagactcgagctaatatttgataggcttttctaaggtcaa |
| Bib1A2 pRSF FP | ttgaccttagaaaagcctatcaaatattagctcgagtctggtaaagaa |
| Bib1A2 pRSF RP | gtgatgatgcggcatggtatatctccttattaaagttaaac |
| Bib1 A2 SSM S64C RP | ccgccaatggcacaaacaggagt |
| Bib1 A2 SSM S64C FP | tggcctcgcattgggtttttgtatctccaaaatttattgttga |
| Bib1 A2 SSM S64A RP | ccgccaatggcacaaacaggagt |
| Bib1 A2 SSM S64A FP | tggcctcgcattgggttttgcaatctccaaaatttattgttga |

Table S3. Codon optimized gene fragments

| Name | Sequence |
|---|---|
| Bib1A1 | atgccacatcaccatcaccaccacatgaaccggaatcaaattattgaagaactcgctgtgaatcaccctgcgggtgcgaagctggttgaggtcagcaaggaagagcttacccgggtgtacgggggtggcgatgttcaagcagaaacttctcctgcctgtgcttccgcaatgctttatctgcaggggttacggtgtctctcggtctgagccggatgcactgctaa |
| Bib1A2 | atgccgcatcatcaccatcaccatatgaatcggaacgaagttattgaagagttagcagtcaatcacccagctggtgctaagttagtggaagtatcaaaagaagaattgacgcgcatttacgcgggggcgatgttcaggcagaaaccactcctgtttgtgccattggcggtggcctcgcattgggtttttcaatctccaaaatttattgttga |
| Bib1Mfrag1 | atgaatcaaaatacaagcataaaaccccttatctctaccactctgtttaaatctctgagtatcaaagaacggattcagcttgcagataaaattgatacgattgacctcaatgaggaggaatttcagagttatcgcaataaatggaaagaagaaacgctgcttaacgatgattccatgaataagagaatacaggtcgagcaactggatgagaaactctttataaaagcgttgaaggcaagtgaaaacgaggaatggcttcaacaaaacaagctggaaccgaaccaaaatccactttggatgaattggattgaggaagcactcacccttcatcgtaataaggagcttaatgaaccagatcaactccatatttcattagcattccgaccgttcatgttatgggctaagaatcgatttaacaaattttatcaggaacacccgtcgctttcaaaaaaagttgattgggatcagttgcagtcttctattctgtataatctgattgaggccctgacaaatattggagcgcg |

| | |
|---|---|
| | cacaatcactttagaattatatattgcaaaacagttaaaagaattaacaggcaacacgccggaag<br>aacgcttcgagtcttttgtgcggactaaacttctggactttgatgcgctggaatttatttataaagaa<br>tatccggttctgtcgcgtattttgatgatacgtactaactattatgtaaacgcaattacagaagccct<br>gacccgttttgaggaagattgggaccagattaaccaaagtttaaaactggatgctttatcgttaaa<br>aagtatcgccgtcggattaggggatagtcatcagcagggtcgaagcgtaatgcgctttaaatttg<br>aaaaaaacaaagagatattatataagccgaaaccattaaccgttgctatttatttccacgagctgc<br>tggattggattaacaaaaagggttttactccaaaattgaaaggccataacattttaaacagacaag<br>gctatgtctgggaagaatgcattcaacatcgagaatgcaaaacccaggaccagattgaaaatta<br>ttacaagcggctcggtgggtatttagctatattaaacgcagtgaatggtactgatttccaccatga<br>aaatatagtcgcggacggcgaatttccgacgctgatcgatttagaaacgatttttcaccatcctgc<br>caagttgaatataaagactgcagaaatcgaggctaaatacaagatcattaattcagtgttaggga<br>ccgctttgcttcctcacttatattttaaaaatgccgagggctatggtattgatatcagcggcgtgag<br>tgtccagaaccaggaggagctgccgattccgttactccgtccggagaacgaggggactgatg<br>agatgcgtttcgtccgcaaaaaagttatggctgatataaacgataacaac |
| Bib1Mfrag2 | gtgccaaagctgaacggcaaagtaattggagcatctagtcacgtggattccatcatcgagggct<br>accagcacgcggcccagataatttttaataataaatccgaactgctgcatgaagacgggcctat<br>agcaaaattcaaggataccgaaattagaattgttgtgcgtcccacacagtactatggcaattttct<br>gcttgaaaaatatcatcctgattacatgcgtgactgcgtggaattagaaaaactgttggatcgact<br>ttggtttaccgtacttgacacgcgacagattccgttcgagaaacaggatttgtttaatggtgatatt<br>cccatttttacaactaagcctggatccagagatctgtttgcctcatccggtgagaaaatagaaaat<br>tattttgatcagccgtcctacgaaatcgtggttgaacgaattaaaaacttaaccctcgattctattga<br>agaacagagtaaatggatcgaggctagcctgagctgcaacattaaagagaaagttattgtgaa<br>ggaagctacatatctcaaagaagaattaaaagaaatcaagaccgatattttcatcgaagaggcc<br>aagaacattggttatcgtctgaaggagcaggttattcatggctcgtataatgatgccacctgggt<br>gggtctcggtatgaattaccacaatcaatggcaagtaactgcactggattcaggattatataacg<br>gtttgtcaggcattgcaatgttcctgggataccttggtaaagtgtctggtgaagaagactttaacc<br>ggttagcaatgcagaccatggaatccatattgcagcaaccgattcaagaaaaaagctttgcatct<br>gcgttttatgggcaggccagtcacctttatatcttatctcattttgacgccctgtatggcgaaaacc<br>aaaaatggaagctgtacattcaaaactctctgaacaacattgaaaagagcgtgaaaaatgacca<br>gttctatgatttattgggcggaagtgccgggattattcaggttctgttaaatatacatgaacagttta<br>ataatgagcaggcgttaaatatagcacaaaaatacggcaatcatttaattgaaaataaaatcgtg<br>acagagagaggtatcgggtggtcagatccttcgagccaaatcatgctcggcggactgagccat<br>ggcacctcgggcatcgcttggtctctgcttcggcttcataagcagaccagcaaacagaagtact<br>ttgaaaccgccctggaggcgatccgatatgaccgtagcctttataacagcaagaactgtaattg<br>ggaagatttaagatgttcccaacataaatcgtctgaattcagtgcagcgtggtgtcatggtgcca<br>ctggcatagggctcagtcgactgttatacctcccctacattaaagatacacttttgcaacaggaga<br>ttgaaaccgcggtcagtactacagtggatgttgggatggggcgcagtcacagtctctgtcatgg<br>tgacctcggtaatagcgaactgtttcatgtggcagggaacgttctcggaagaccagagtggaat<br>agaatggcgcacgcggtaggtatgaacaccataaaagagaaacaagaaattggaaagtacaa<br>aacaggcgtaggtcgtcatatcgagatccctggactgtttatgggcctgtcaggaatcggatac<br>cagctcttacgtctggctaagccaaaccaagttcccagtgtcttgaccttagaaaagcctatcaa<br>atattag |

Table S4. The accession identification numbers associated with the corresponding proteins named in this work

| Accession | Name |
| --- | --- |
| WP_098353184.1 | Bib1R1 |
| WP_000933887.1 | Bib1R2 |
| WP_098353186.1 | Bib1M |
| WP_016078320.1 | Bib1 A1 |
| WP_016078319.1 | Bib1 A2 |
| WP_098353188.1 | Bib1A |
| WP_098353190 | Bib1B |
| WP_016078316.1 | Bib1R |

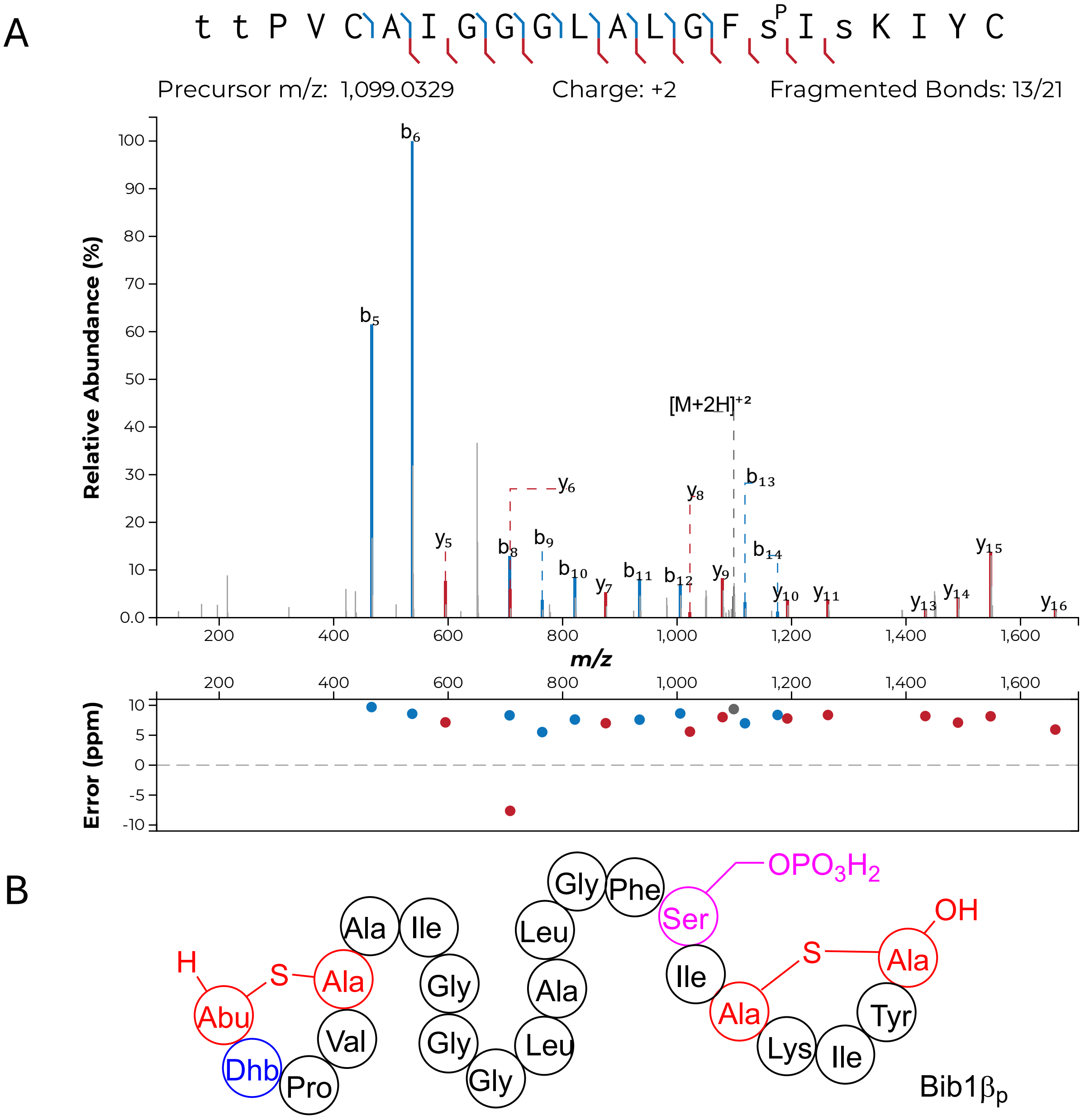


Fig. S1. LCMS-MS analysis and ring pattern of Bib1βP. A) Analysis of the fragmentation pattern produced during tandem MS-MS. Lower-case labels indicate that a residue was post translationally modified. '$s^P$' indicates that a Ser was phosphorylated. All other lower-case residues were dehydrated. B) Ring pattern and phosphorylation site consistent with the LCMS-MS analysis.

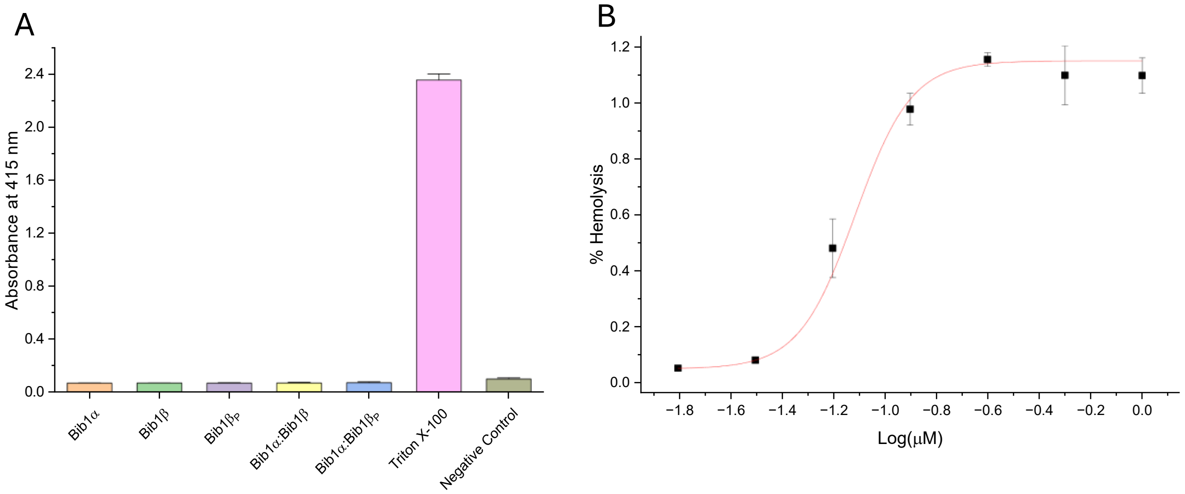


Fig. S2. Bibacillin 1 does not show hemolytic activity in solution at ≤2 μM. A) Hemoglobin release (measured by absorbance at 415 nm) after treating rabbit erythrocytes with Bib1α, Bib1β or Bib1$\beta_P$ individually or with a 1:1 of Bib1α:Bib1β or Bib1α:Bib1$\beta_P$ for 1 h at 37 ℃. As a positive control, erythrocytes were treated with 0.03% v/v triton X-100. Negative control contained only buffer. A total peptide concentration of 2 μM was used in each case. B) Extent of hemolysis after treating freshly washed rabbit erythrocytes with 1:1 of CylL$_S$”:CylL$_L$” at 37 ℃ for 1 h. Data was fit to a dose response curve using Origin 7 yielding an $EC_{50}$ = 76 ± 5 nM.

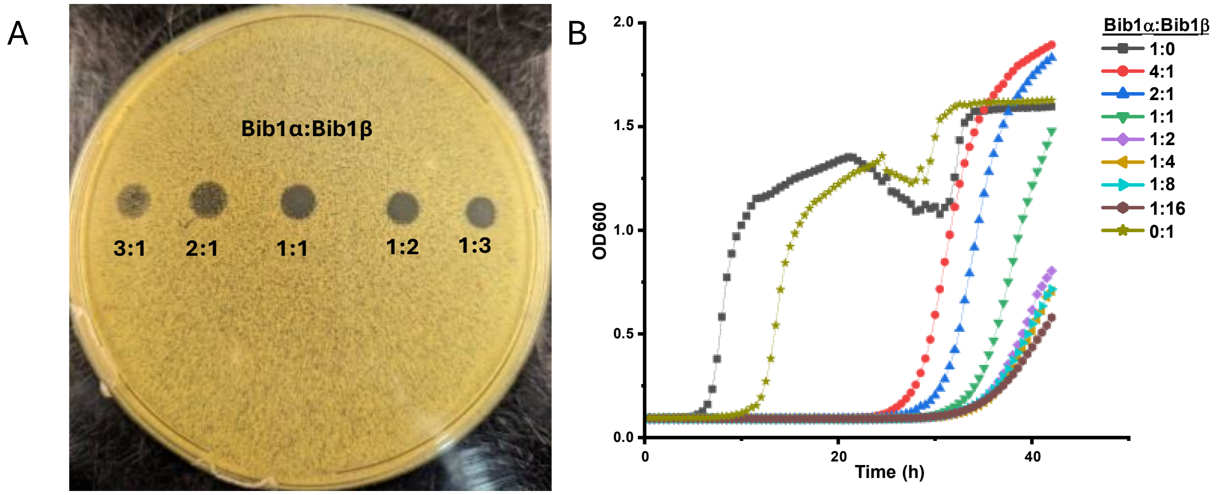


Fig. S3. Continuous variation experiments demonstrate that bibacillin 1 does not produce maximal activity at a well-defined ratio of Bib1α and Bib1β. A) Agar diffusion assay using *Micrococcus luteus* ATCC 4698 seeded soft agar. Bib1α and Bib1β were diluted to 100 μM in PBS then mixed together to produce the indicated ratios and 3 μL of each mixture was spotted on the agar plate. The total amount of peptide deposited was 300 pmol in each case. B) Growth curves collected simultaneously with those featured in Figure 4E. In this case, the total peptide concentration was kept at 5 μM.

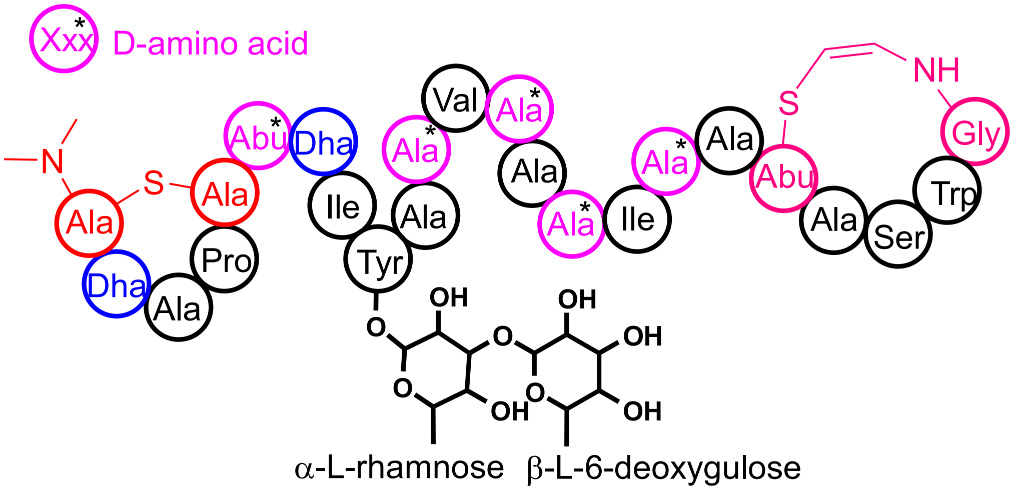


Fig. S4. The structure of cacaoidin. Post translationally modified residues are color coded. Red residues comprise an *N*,*N*-dimethylated lanthionine. Blue residues are dehydrated amino acids. Magenta residues containing a black asterisk are D-amino acids. Residues involved in S-[(Z)]-2-aminovinyl- (3S)-3-methyl]-D-cysteine are colored rose pink.

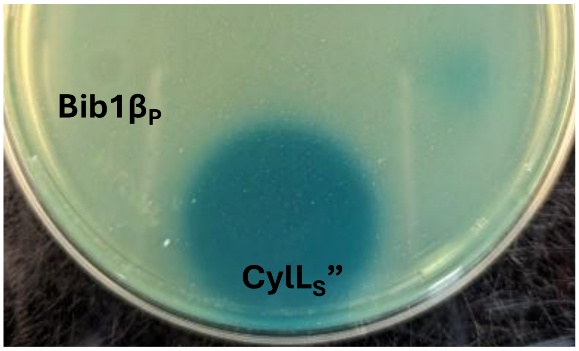


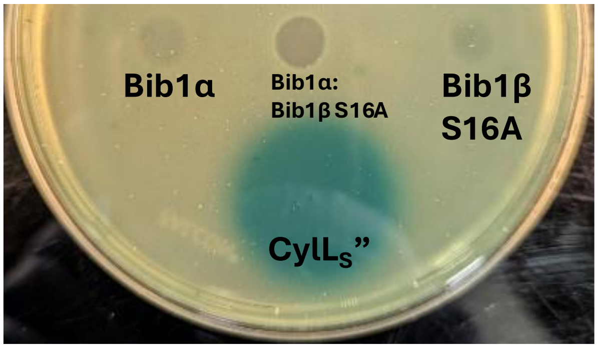


Fig. S5. Bib1βP and Bib1β S16A do not trigger the CylR1/R2 system. Plates were prepared as described in Figure 6A.

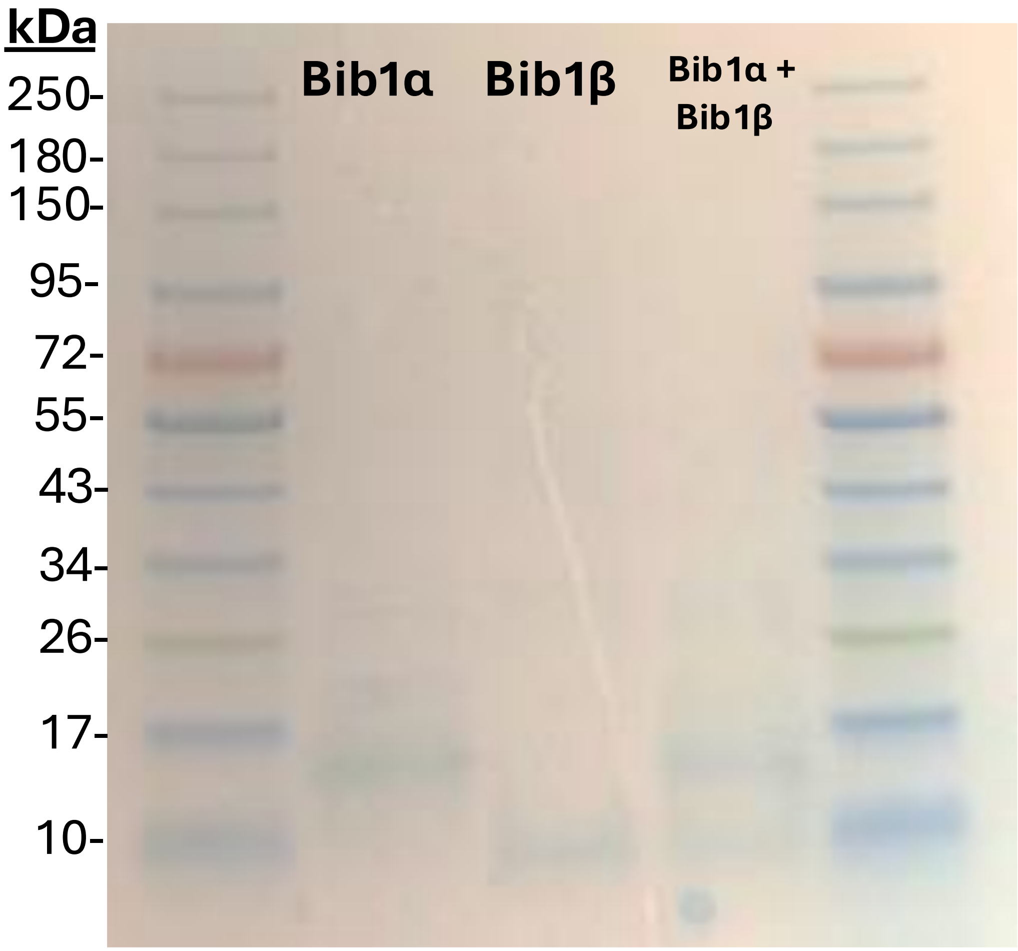


Fig. S6. SDS-PAGE of Bib1α, Bib1β or Bib1α:Bib1β (1:1). In total, 2 nmol of peptide was present in each lane. A gradient of 4-20% acrylamide concentration was used. Coomassie staining was used to image peptide species.

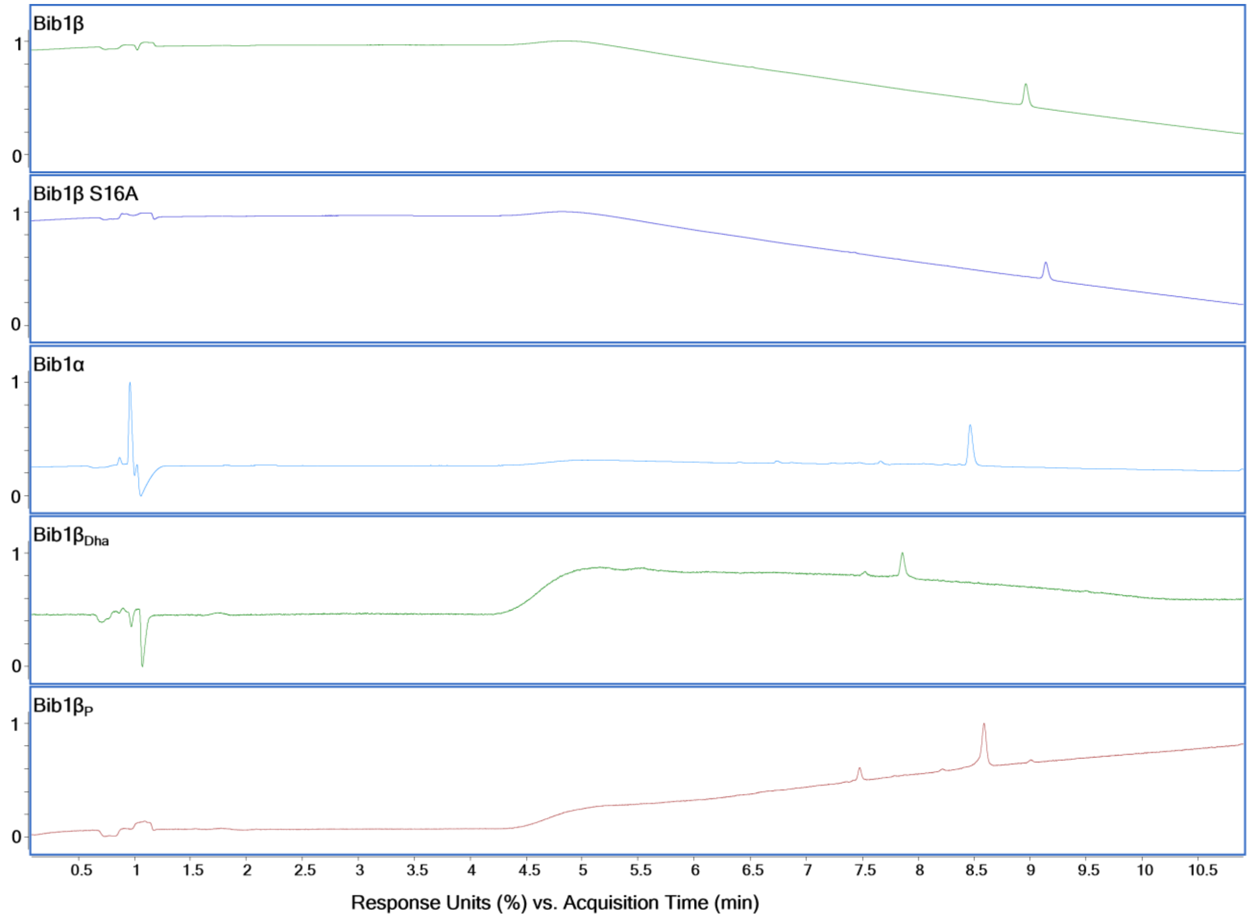


Fig. S7. Reversed phase-HPLC traces of purified Bib1α, Bib1β, Bib1$\beta_P$, Bib1$\beta_{Dha}$, and Bib1β S16A. Traces were collected using the conditions described in the High-Resolution Tandem Mass Spectrometry section of the Experimental.

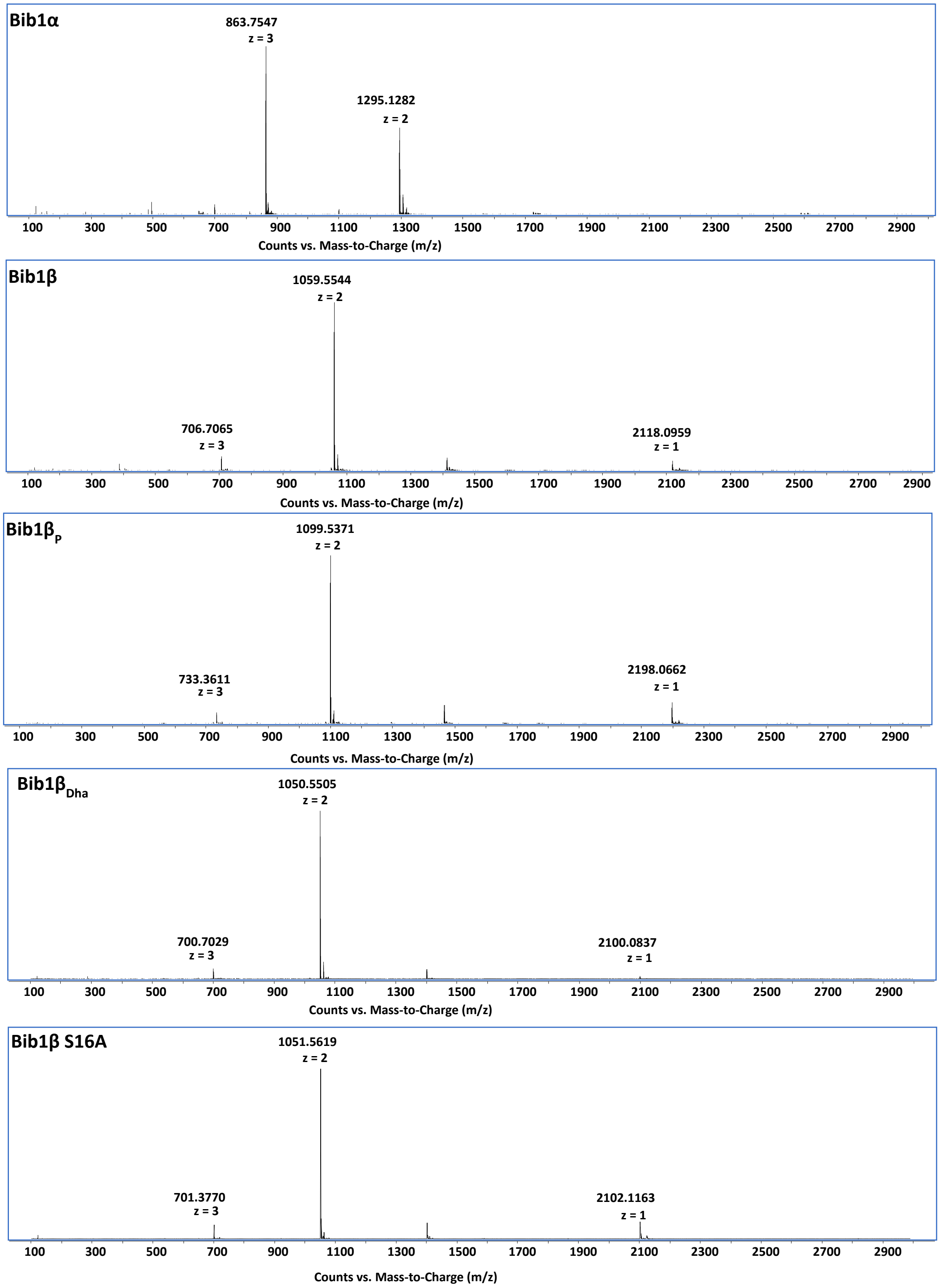


Fig. S8. HRMS spectra of purified Bib1α, Bib1β, Bib1β$_P$, Bib1β$_{Dha}$, and Bib1β S16A. HRMS data was collected using the conditions described in the High-Resolution Tandem Mass Spectrometry section of the Experimental. For calculated m/z values, see Table S5.

Table S5. Comparing high resolution m/z to expected m/z for Bib1α, Bib1β, Bib1$β_P$, Bib1$β_{Dha}$, and Bib1β S16A

| Peptide | Expected m/z $[M+2H]^{2+}$ | Observed m/z $[M+2H]^{2+}$ | Δ (ppm) |
|---|---|---|---|
| Bib1α | 1294.6233 | 1294.6255 | 2.2 |
| Bib1β | 1059.0497 | 1059.0559 | 6.2 |
| Bib1$β_P$ | 1099.0329 | 1099.0344 | 1.5 |
| Bib1$β_{Dha}$ | 1050.0445 | 1050.0485 | 4.0 |
| Bib1β S16A | 1051.0523 | 1051.0603 | 8.0 |
